# Mapping Endothelial-Macrophage Interactions in Diabetic Vasculature: Role of TREM2 in Vascular Inflammation and Ischemic Response

**DOI:** 10.1101/2024.05.14.594235

**Authors:** Naseeb Kaur Malhi, Yingjun Luo, Xiaofang Tang, Rahuljeet Singh Chadha, Alonso Tapia, Dongqiang Yuan, Shufan Yin, Xuejing Liu, Muxi Chen, Meirigeng Qi, Marpadga A. Reddy, Lu Wei, John P. Cooke, Esak Lee, Rama Natarajan, Kevin W. Southerland, Zhen Bouman Chen

## Abstract

Diabetes mellitus (DM) significantly accelerates vascular diseases like peripheral arterial disease (PAD). Endothelial cells (ECs) and macrophages (MΦs) singularly and synergistically are important contributors to DM-associated vascular dysfunction. Single-cell (sc) profiling technologies are revealing the true heterogeneity of ECs and MΦs, but how this cellular diversity translates to cell-cell interactions, and consequentially vascular function, remains unknown. We leveraged scRNA sequencing and spatial transcriptome (ST) profiling to analyze human mesenteric arteries from non-diabetic (ND) and type 2 diabetic (T2D) donors. We generated a transcriptome and interactome map encompassing the major arterial cells and highlighted Triggering Receptor Expressed on Myeloid Cells 2 (*TREM2*) as a top T2D-induced gene in mononuclear phagocytes (MPs), with concomitant increases of TREM2 ligands in ECs. We verified DM-associated TREM2 induction in cell and mouse models, and found that TREM2 inhibition decreases pro-inflammatory responses in MPs and ECs, as well as increases EC migration *in vitro*. Furthermore, TREM2 inhibition using a neutralizing antibody enhanced ischemic recovery and flow reperfusion in DM mice subjected to hindlimb ischemia, suggesting that TREM2 promotes ischemic injury in DM. Finally, in human PAD, co-existing DM was associated with greater expression of TREM2 and its interaction with ECs, with a further increase in ischemic tissue compared to patient-matched non-ischemic tissue. Collectively, our study presents the first atlas of human diabetic vessels with single cell and spatial resolution, and identifies TREM2-EC interaction as a key driver of diabetic vasculopathies, the targeting of which may offer an opportunity to ameliorate vascular dysfunction associated with DM-PAD.

## Introduction

Diabetes mellitus (DM) is a major global epidemic negatively affecting over half a billion adults worldwide, creating a substantial healthcare burden (1). DM predisposes and accelerates various vasculopathies including, but not limited to, coronary and peripheral arterial diseases (PAD). The latter restricts perfusion to the lower limbs and is nearly four times more prevalent in individuals with DM; yet there are very limited therapeutic options. Patients with DM-PAD are also more likely to progress to critical limb-threatening ischemia (CLTI) (2). While it is well known that DM can cause chronic inflammation and tissue injury (3–5), the mechanisms underlying DM-induced vascular dysfunction, remain incompletely understood, which hampers the development of effective therapeutics for managing diabetic vasculopathy.

As with any given disease, diabetic vasculopathy involves a multitude of cell types and complex cellular communications and interactions. One of the earliest pathogenic events to occur in DM is endotheliopathy, i.e., the dysfunction of endothelial cells (ECs) lining the vascular lumen. Apart from the well-recognized role of EC dysfunction as a result of DM and a driver of diabetic vasculopathy, recent studies have also highlighted EC dysfunction in the development of metabolic dysfunction and DM per se (6–8). While quiescent ECs express low levels of adhesion molecules (e.g., vascular cell adhesion molecule 1 [VCAM1] and intercellular cell adhesion molecule 1 [ICAM1]), the diabetic milieu (e.g., hyperglycemia and elevated TNF-α levels) cause ECs to become more adhesive, enabling them to recruit monocytes, which adhere and transmigrate into the sub-endothelial layer and differentiate into macrophages (MФs) (9). The differentiated MФs can further propagate the inflammatory response and vascular dysfunction (3).

While this classical diapedesis and pro-inflammatory process involving ECs and monocytes/MФs (collectively termed mononuclear phagocytes, MPs) in diabetic vasculopathy as well as other vascular diseases has been well recognized, the EC-MP interactions have mostly been studied in a unidirectional fashion. New insights have unraveled bi-directional communications between ECs and MPs. For example, patrolling monocytes crawling along the vascular endothelium can monitor ECs and remove the damaged ECs (10–12). Intimal resident MΦ are essential for maintaining a non-thrombogenic endothelial state (13, 14). Under hypoxia or ischemia, MΦ can modulate angiogenesis and collateral arterial remodeling, possibly through vascular endothelial growth factor (VEGF) and monocyte chemoattractant protein-1 (MCP-1) signaling (15–17). In cancer, activated MΦ can secrete inflammatory mediators to disrupt EC tight junctions and increase EC permeability (18). Furthermore, recent studies leveraging single-cell (sc) technologies have revealed unexpected heterogeneity of MΦ (19–22), with various subtypes featuring distinct gene expression profiles and associated with tissue-damaging or -reparative properties (13, 23). Despite the known complexity of EC-MΦ interactions, the specific ligands and receptors driving the diverse EC-MΦ interplay —and their varied functional outcomes—remain largely uncharacterized, underscoring a critical gap in our understanding of vascular inflammation and dysfunction in diabetic contexts.

In this study, we aim to identify key EC-MФ interactions contributing to diabetic vasculopathies, particularly PAD. We began with scRNA and spatial transcriptome (ST) profiling of human mesenteric arteries from non-diabetic (ND) donors and donors with type 2 DM (T2D) and mapped the DM-associated EC-MΦ interactions in the arterial wall. Through unbiased analyses, we identified Triggering Receptor Expressed on Myeloid Cells 2 (TREM2), a phagocytic cell-surface receptor, to be highly upregulated in MPs in the diabetic intima, concomitant with its interaction with dysfunctional ECs in DM. The DM-induced increase of TREM2 in MPs was validated by *in vitro* and mouse models of DM. Importantly, we identified a novel role of TREM2 in promoting DM-induced vascular inflammation and impairing EC wound-healing capacities. In DM mice with hindlimb ischemia (HLI), blocking of TREM2 using a neutralizing antibody (TREM2-Ab) enhanced ischemic recovery and flow perfusion, suggesting a role of TREM2 in promoting diabetic PAD (DM-PAD). To provide the translational relevance of our findings, we demonstrate an increase of TREM2 and its interaction with ECs in human patients with DM-PAD. Collectively, our study presents an atlas of human diabetic vessels with a focus on EC-MP interactions. Exemplified by TREM2-EC interactions, our findings provide valuable new insights into EC dysfunction and EC-MΦ interactions, key processes contributing to diabetic vasculopathies such as PAD.

## Results

### Profiling the diabetic transcriptome in human mesenteric intima

We began by profiling the DM-associated transcriptomic changes in vascular cells, through scRNA-seq of human mesenteric arteries from 10 deceased organ donors with or without DM (5 ND and 5 T2D) collected between 2019 and 2024, a tissue source we have previously described (24–29) (**Table S1, fig. S1**). The reasons for using mesenteric arteries include tissue availability and the demonstrated utility of using these vessels to study DM-induced vascular dysfunction (30–32). The 5 ND donors included 1 Hispanic and 4 Caucasians, all men. The 5 T2D donors consisted of 4 Hispanic individuals (1 woman and 3 men) and 1 Caucasian man, with 4 of the donors classified as obese (BMI≥30). Of note, all four Hispanic donors were untreated for their T2D, and the one Caucasian donor was noncompliant with treatment.

Based on the existing literature reporting scRNA-seq of whole blood vessels, ECs represent only 5-15% of the entire population (33–37). To enrich ECs and capture EC-interacting cells (esp. MΦ), we isolated the inner layer of the arterial wall (including intima), using an intima scRNA-seq protocol we have described (25) (**Fig. 1A**). From 10 donors, we retrieved ∼14,000 cells in total, all of which passed quality control for scRNA-seq. Principal component analysis (PCA) of the batch-corrected intima scRNA-seq data revealed slight separation between ND and T2D, implying generally similar cell populations obtained from ND and T2D donors (**Fig. 1B**). Based on unsupervised clustering, these cells were separated into 21 clusters (**fig. S2**). We chose to use unbiased, default methods of analyses, to reduce the possibility of over-interpretation of our scRNA-seq data. As a result, monocytes (in potentially differentiating states) and MΦ were assigned collectively as mononuclear phagocytes (MPs). Annotation of all intima scRNA-seq cells using reported cell type marker genes (34, 35, 37) identified 6 major cell types: ECs (∼34%, consisting of 2 clusters), MPs (∼27%, consisting of 5 clusters), T/NK cells (∼30%, consisting of 6 clusters), VSMC/fibroblasts 2%, consisting of 2 clusters), and others (e.g., B cells) (**Fig. 1C, D**). This cell composition is expected considering the intima dissociation and EC enrichment.

**Figure 1.**
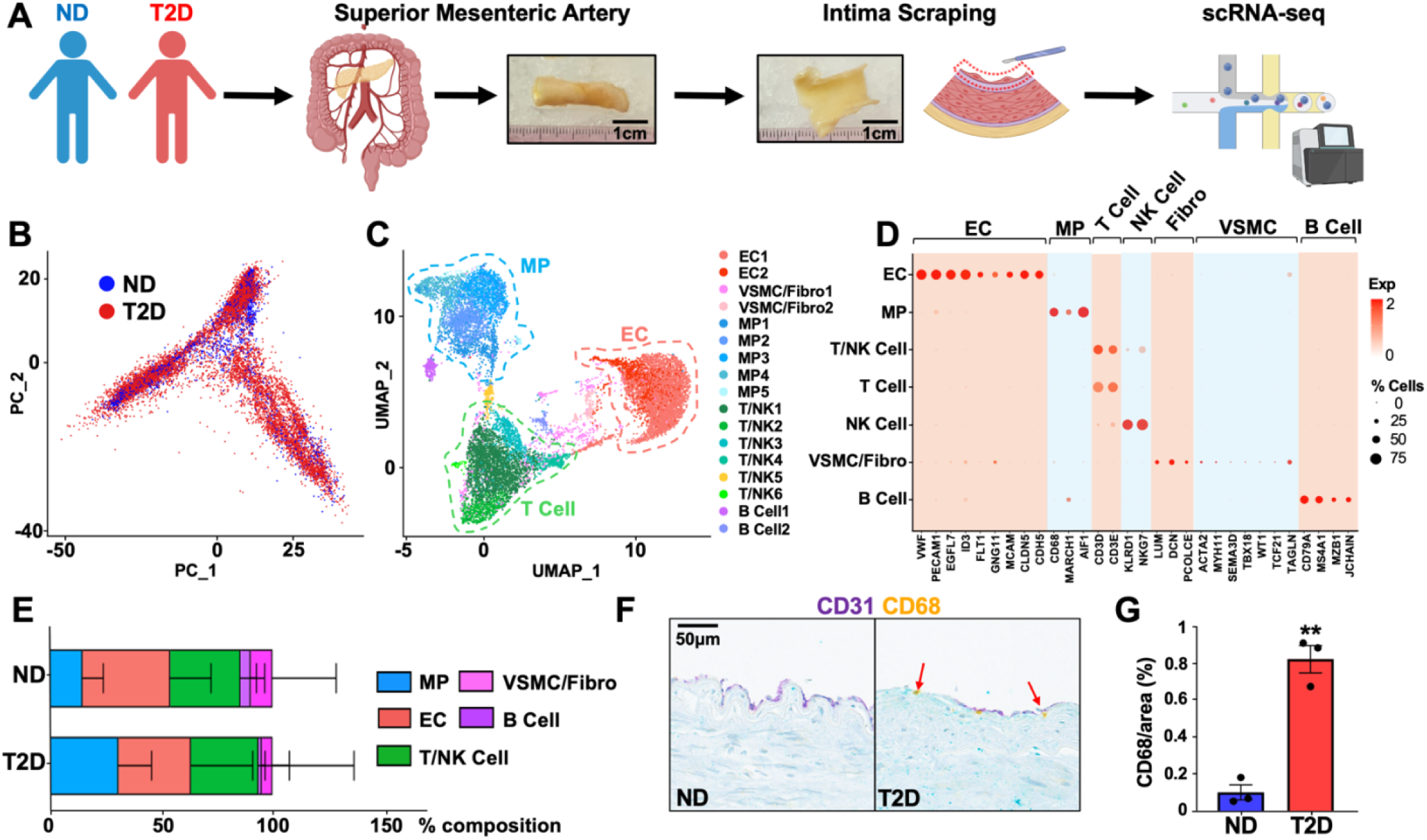
Profiling of human mesenteric artery intima. **(A)** Schematic of human mesenteric artery processing for scRNA-seq. ND: Non-diabetic; T2D: Type 2 diabetic. **(B)** Principal component analysis (PCA) plot of intima scRNA-seq data, sorted by diabetic state. **(C)** Uniform manifold approximation and projection (UMAP) of scRNA-seq data, classified by major cell types and subclusters generated through Seurat. EC: endothelial cell; VSMC: vascular smooth muscle cell; MP: mononuclear phagocyte; NK: natural killer. **(D)** Dotplot showing levels of cell-marker genes (x-axis) within each annotated cell population (y-axis). **(E)** Cell composition (%) of the intima scRNA-seq data from ND vs T2D donors. **(F, G)** Representative CD31 and CD68 co-staining and CD68 quantification of mesenteric arteries (n=3 ND/T2D). Red arrows indicate CD68-positive staining. Bar graphs represent mean±SEM. **p<0.01 compared to ND based on Student’s t-test.

Comparing the cell composition between ND and T2D, we observed no major difference in expression of cell marker genes in most clusters (**fig. S2**). When we compared the distribution of the major cell types, while the percentages of ECs and T cells were not substantially different (39 to 32% for ECs and 31 to 30% for T cells), that of MPs increased from 15% in ND to 31% in T2D (**Fig. 1E**). We validated this at the protein level, observing more intense CD68 (an MP marker) staining in T2D intima as compared to ND (**Fig. 1F, G**). This is in line with the notion that DM induces inflammatory infiltration and MΦ accumulation in the sub-endothelial layer (38, 39).

### Endothelial and macrophage diversity in the diabetic intima

We next investigated the DM-associated transcriptomic changes within ECs and MPs. We first focused on ECs. Two EC clusters were identified from the intima scRNA-seq, EC1 (79% of ECs) and EC2 (21% of ECs) (**Table S2**). The proportion of EC clusters did not differ significantly between ND vs T2D. While EC1 expresses higher levels of genes encoding extracellular matrix components (e.g., *COL8A1*, *ELN*), EC2 expresses higher levels of genes related to lipid metabolism (e.g., *FABP4* and *FABP5*) (**fig. S3**), similar to EC clusters previously described (33, 40, 41). We then analyzed DEGs by comparing T2D- vs ND-EC data using RePACT, a regression-based algorithm that has shown improved sensitivity in identifying disease relevant genes even with a small number of samples (42, 43). RePACT analysis revealed a remarkable difference in the transcriptome states between ND- vs T2D-ECs (**Fig. 2A-C**), with DEGs significantly enriched for wound healing/angiogenesis, oxidative stress and inflammatory responses **(fig. S4A, B)**. For example, genes promoting inflammatory response and endothelial-mesenchymal transition (e.g., *CCN2*, *THBS1*, and *FN1*) are induced, whereas genes crucial for EC homeostasis and angiogenesis (e.g., *NOS3, NRP1, and PECAM1*) are suppressed in T2D-ECs (**Fig. 2C, fig. S4A, Table S3**). These changes are in line with the previously reported effect of DM conditions on ECs (44), which also validates the use of these human data to investigate DM-induced vascular changes.

**Figure 2.**
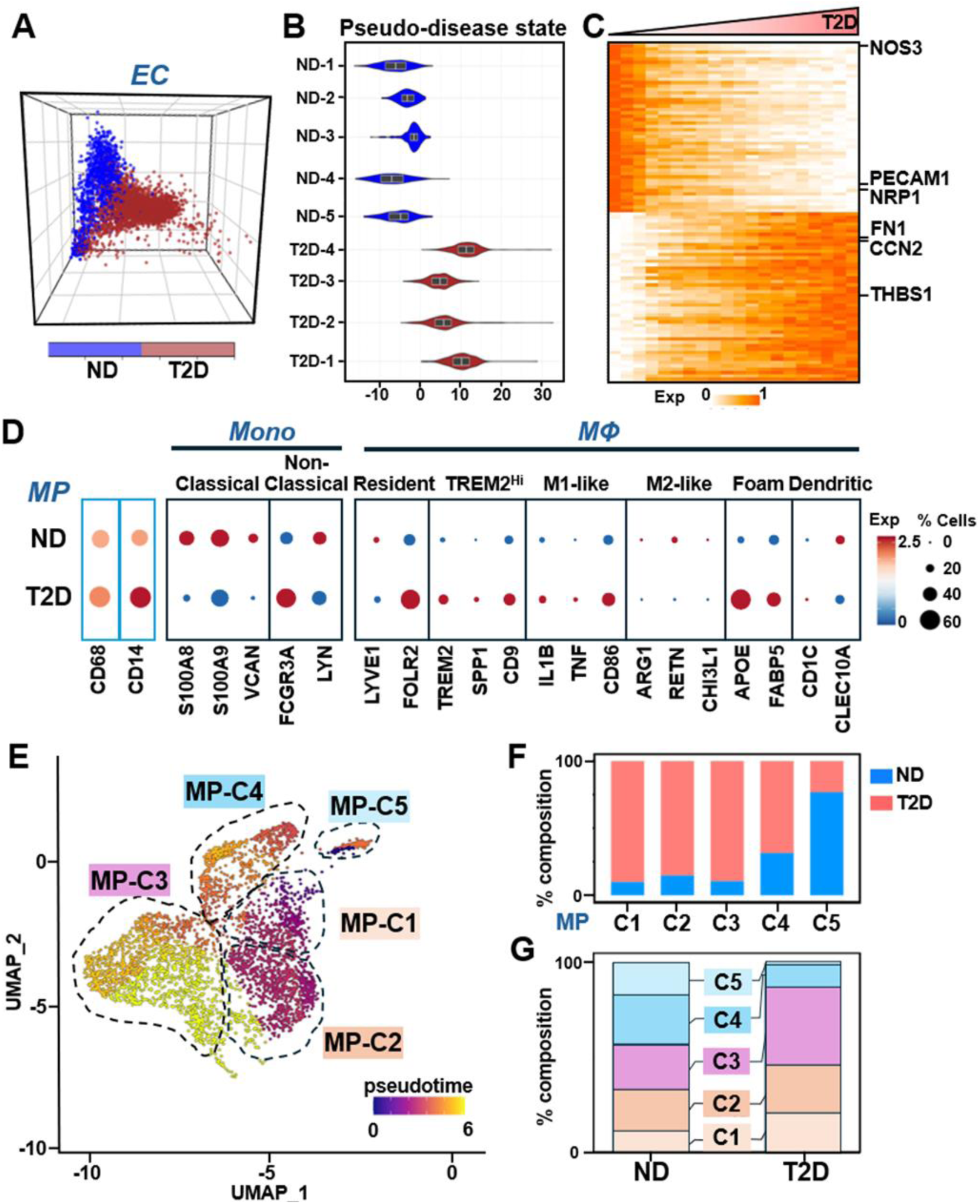
DM-induced transcriptome shifts in EC and MP. (A-C) MP profiled by intima scRNA-seq. **(A)**. ECs from ND (blue) or T2D (red) vessels are plotted in the 3D space of the top 3 principal components. **(B)** Violin plots showing the pseudo-disease state in accordance with RePACT analysis of each donor. **(C)** Heatmap showing top 50 up- and down-regulated DEGs in T2D vs ND ECs. Each row represents the expression of a DEG in single cells ranked by the pseudo-disease state. **(D-G)** MP profiled by intima scRNA-seq. **(D)** Expression of subtype markers in MP of ND and T2D plotted with z scores. **(E)** MPs colored by pseudotime as calculated by Monocle 3 superimposed on UMAP. Note that MP-C3 progressed furthest through pseudotime. **(F)** Contribution of ND vs T2D samples to each of the five MP clusters (MP-C1 toC5). **(G)** Percentage of MP-C1 to -C5 in total MP population from ND and T2D samples.

We analyzed in parallel the DM-associated MP transcriptome and identified 935 DM-induced DEGs, including 473 up- and 462 down-regulated (**fig. S5A, B**). The top enriched pathways are involved in immune and inflammatory responses, as well as angiogenesis (**fig. S5C**). Given the heterogeneity of MPs, we also examined expression of marker genes of several well-known myeloid subtypes within the MP population (22, 37, 45). The common MP marker genes, i.e., *CD14* (for monocytes) and *CD68* (for MΦ) showed a slight but insignificant increase between ND and T2D (**Fig. 2D**). There was an apparent decrease in classical monocyte markers, e.g., *S100A8*, and *VCAN* in T2D, suggesting more differentiation into MΦ. While the M1-like MΦ markers, e.g., *IL1B* and *TNF*, were increased, the M2- like markers, e.g., *ARG1* and *RETN*, were decreased, indicative of a more pro-inflammatory state of MΦ under DM conditions. Interestingly, the marker genes of foam cells (i.e., *FABP5* and *APOE*) and TREM2^Hi^ MΦ (an important subtype implicated in many diseases including DM marked by *TREM2*, *SPP1* and *CD9*) (46–50), were also increased in T2D MPs (**Fig. 2D**).

Next, we performed trajectory analysis using Monocle 3 to infer DM-associated cell state changes (51), focusing on ECs and MPs. While ECs showed no obvious movement through pseudotime (**fig. S6**), the MP population, consisting of five clusters (hereafter named MP-C1 to C5), exhibited a clear trajectory. Specifically, MP-C3 showed the furthest progression through pseudotime (**Fig. 2E**), suggesting that this cluster represents the most advanced T2D state. While MP-C5 is more dominant in ND samples, MP-C1 to C4 are more present in T2D. Among these, MP-C3 demonstrated the largest percentage increase in MPs in T2D vs ND (**Fig. 2F, G**), suggesting an important role of this cluster in DM-induced vascular inflammation and dysfunction. Pathway enrichment analysis of DM-associated changes in the MP clusters revealed that MP-C3 was enriched in genes involved in angiogenesis, hypoxia response, and NO biosynthetic process, all EC-relevant pathways, and pathways regulating inflammatory response and cytokine production (**fig. S7, 8**).

We then sought to determine the key genes within the MP-C3 contributing to the T2D state. To this end, we identified 65 genes (46 increased and 19 decreased) that were consistently differentially expressed between ND vs T2D in all the MP population as well as in MP-C3 (**fig. S9**). Among these genes, the top ranked is *ADAM28*, a metalloprotease with functions in cell adhesion, migration, proteolysis, and signaling (52). The second ranked gene is *TREM2*, a promiscuous receptor that has been shown to primarily express on the surface of myeloid cells and regulate the immune response (47). Other notable genes include Histamine Receptor H2 (*HRH2*) and plasminogen activator, urokinase (*PLAU*), to which antagonists have been developed for potential DM treatment (53, 54) (**fig. S9**).

Additionally, we identified significant changes in the T cell, NK cell and B cell transcriptome between ND and T2D (with 425, 325, 862 DEGs respectively) (**fig. S10**). Given the focus of the current study on ECs and MΦ, we did not perform further analyses of these cells.

### Increased EC-MP interactions in T2D mesenteric artery

To identify EC-MP interactions with a spatial context, we performed spatial transcriptome-seq (ST-seq) using 10X Genomics Visium platform, which captures RNA on a tissue slide (6.5 mm x 6.5 mm) consisting of ∼5,000 spots, termed voxels (50 μm diameter) with embedded spatial barcodes. We performed ST-seq on 4 mesenteric arteries (2 ND vs 2 T2D) (**Table S1**). These were different donors from those used for intima scRNA-seq due to the limited tissue size. While the ND arteries showed no obvious pathologies, the DM arteries show intima-media thickening (**fig. S11**). Through the Visium workflow, we captured 2,000 UMI counts from 60,000 reads per voxel and 17,000 genes per sample (**fig. S12**).

We first anchored the ST-seq data based on histology. As expected, *VWF* (an EC marker gene) was detected in the intima, where *CD68* was also detected (**Fig. 3A**). These data indicate that the ST-seq data indeed retains the spatial features of the gene expression and captures EC and MΦ in proximity. Given the 50 µm resolution of our ST-seq, which will not provide gene expression profile at single cell levels (**fig. S13A, B**), we deconvoluted the ST-seq data using scRNA-seq data. For this purpose, we performed scRNA-seq of the whole arteries, which have cellular composition matching that of ST-seq to enable efficient data integration and spatial anchoring (**fig. S13C-F**). Through the integrative analysis of ST- and scRNA-seq, each voxel was assigned probability scores for one of the five major cell types in the arterial wall (i.e., ECs, VSMCs, MΦs, Fibroblasts, and T cells). Thus, voxels with probability scores greater than 20% for ECs were defined as “EC-dominant voxels”. We then took these voxels and queried information on other present cells in EC proximity (i.e. within 50 µm distance), thus encompassing potential intercellular interactions. While in the ND arteries, the probability of MΦs in the ‘EC-dominant’ voxels was 16%, that in the T2D arteries was 41% (**Fig. 3B**), suggesting an increase in EC-MΦ interactions in the T2D arteries.

**Figure 3.**
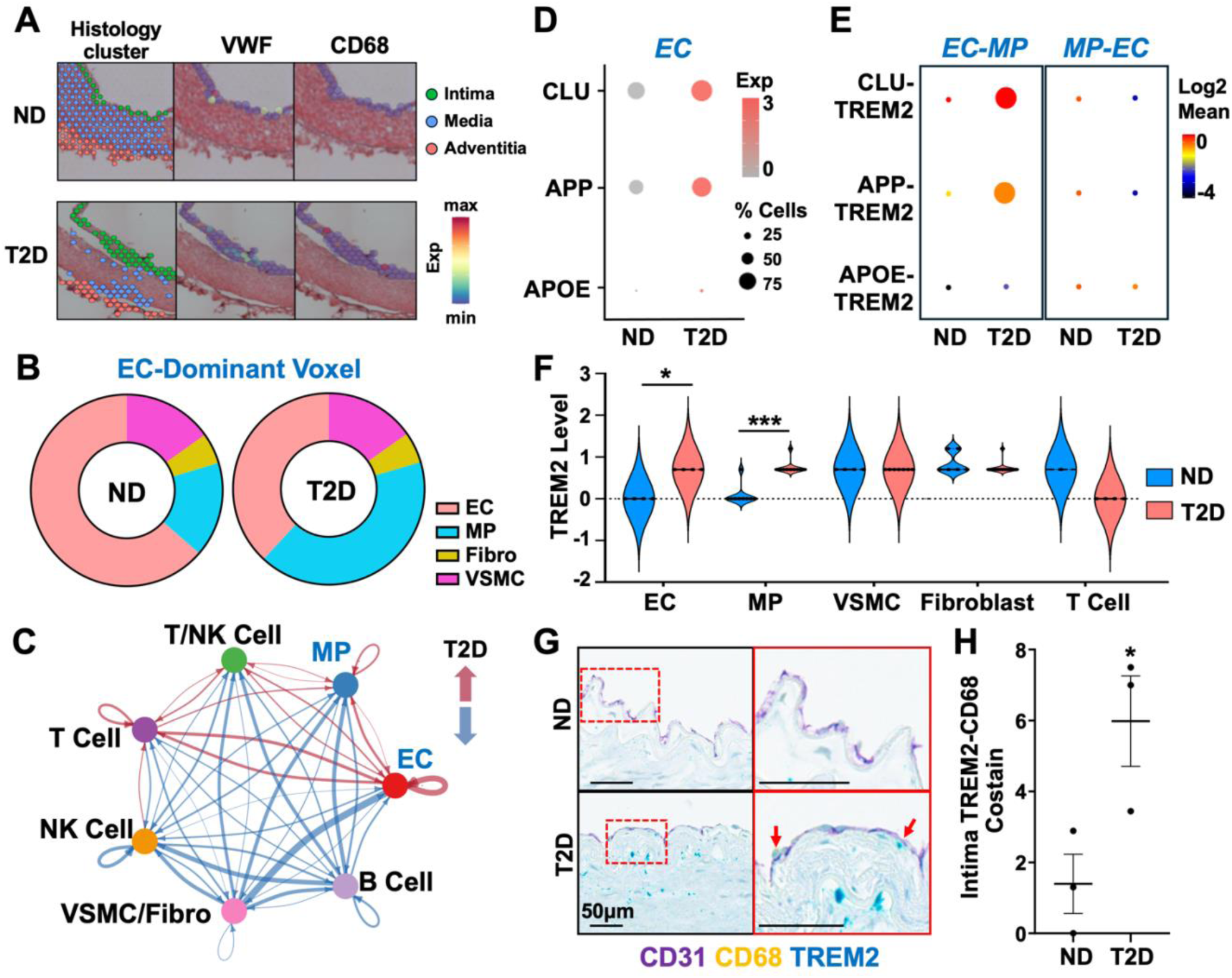
Increased EC-MP and EC-TREM2 interactions in T2D arteries. **(A)** SpatialFeaturePlots showing histology guided annotation (left) used to subset out the intima. Middle and right plots show VWF and CD68 expression in the intima as determined by ST-seq. **(B)** Pie charts showing probability of the presence of specific cell types within ‘EC-dominant’ voxels in ND and T2D mesenteric arteries, computed through integrating ST-seq and whole-artery scRNA-seq. **(C-E)** Analysis for cell interactions based on intima scRNA-seq data. **(C)** Comparison of intercellular interactions between ND vs T2D analyzed by CellChat. Arrows denote intercellular interactions with red indicating increase and blue indicating decrease in T2D as compared to ND. **(D,E)** Gene expression of TREM2 ligands in ND vs T2D ECs (in D) and CellphoneDB analysis of bi-directional ligand-TREM2 pairs between MPs and ECs in ND and T2D arteries (in E) determined by intima scRNA-seq. **(F)** TREM2 expression levels in ST-seq voxels with specific dominant cell types as indicated, determined through integrative analysis of ST- and scRNA-seq. **(G)** Immunohistochemistry of CD31, CD68, and TREM2, from left to right showing increased magnification and zoomed-in images. Red arrows indicate CD68-TREM2 co-positive spots. **(H)** Quantification of CD68-TREM2 co-staining in the intima. *p<0.05, ***p<0.001 based on Student’s t-test.

To validate that the increased EC-MP interactions in T2D arteries occur in the intima and to identify ligands and receptors mediating these interactions, we performed CellChat and CellPhoneDB analyses with intima scRNA-seq data as shown in **Figs. 1** and **2**. These bioinformatics tools allowed inference of cell-cell communication based on gene expression level of ligand-receptor pairs (55, 56). Our analysis revealed an overall increase in intercellular interactions in T2D as compared to ND, evidenced by both the number of interactions and the interaction strength (**fig. S14A**). EC interactions with MPs as well as with other immune (T/NK) cells were all increased in T2D (**Fig. 3C, fig. S14B**). Focusing on the EC-MP interactions, we identified a substantial increase in bidirectional communications, exemplified by EC-derived THBS1, TGF-β, and VCAM pathways and MP-derived CXCL, ICAM, and CSF pathways (**fig. S15A, B**).

Given the emerging importance of TREM2 and TREM2^Hi^-MΦ in various diseases (46–50) and its prominent induction in the diabetic artery intima (**Fig. 2, fig. S9**), we were interested in the role of TREM2 in the DM-induced vasculopathy, which has not been reported. Therefore, we focused on TREM2 as an example to validate the EC-MP interaction and investigate the functional importance of EC-TREM2 interactions in diabetes-associated vascular function.

Consistent with the increased *TREM2* expression in T2D MPs, the expression levels of several known TREM2 ligands, including clusterin (*CLU*), amyloid-β precursor protein (*APP),* apolipoprotein E (*APOE*) (47, 57) were all increased in T2D ECs (**Fig. 3D, fig. S15C**). Moreover, the inferred ligand-TREM2 receptor pairs, i.e., CLU-TREM2 and APP-TREM2 were also increased in DM than in ND (**Fig. 3E**). In line with the scRNA-seq data, the deconvolved ST-seq data revealed that TREM2 expression was increased in not only ‘MP-dominant’ voxels (as expected), but also ‘EC-dominant’ voxels in T2D arteries (**Fig. 3F**). As a validation of EC-TREM2 MΦ interactions at the protein level, in the CD31-stained intima layer, TREM2-CD68 co-stain was increased in T2D vessels (**Fig. 3G, H**). These spatially resolved data, together with the scRNA-seq data, suggest that TREM2-expressing MΦs interact with ECs, especially under DM conditions.

### Diabetic conditions increase TREM2 expression *in vitro* and *in vivo*

As shown in **Fig. 4A, B**, TREM2 expression is clearly increased in the MPs in the human T2D mesenteric artery intima and is highest in MP-C3. To verify the increase of TREM2 expression in the diabetic vasculature, we found that *Trem2* was also increased in the aortas of C57Bl/6 mice treated with streptozotocin (STZ), a model of hyperglycemia and DM (**Fig. 4C**). Moreover, we demonstrated increased *TREM2* expression by combined high glucose and TNF-α (HT, previously used to mimic a DM condition *in vitro*) (24, 58) in human CD14^+^ MPs, THP-1-derived human MФs, and mouse RAW 264.7 MФs (**Fig. 4D, E, fig. S16**). Of note, the increased TREM2 expression levels were attendant with an increase of phospho-Syk, indicative of an increase in TREM2 activity (59) (**Fig. 4F, G**).

**Figure 4.**
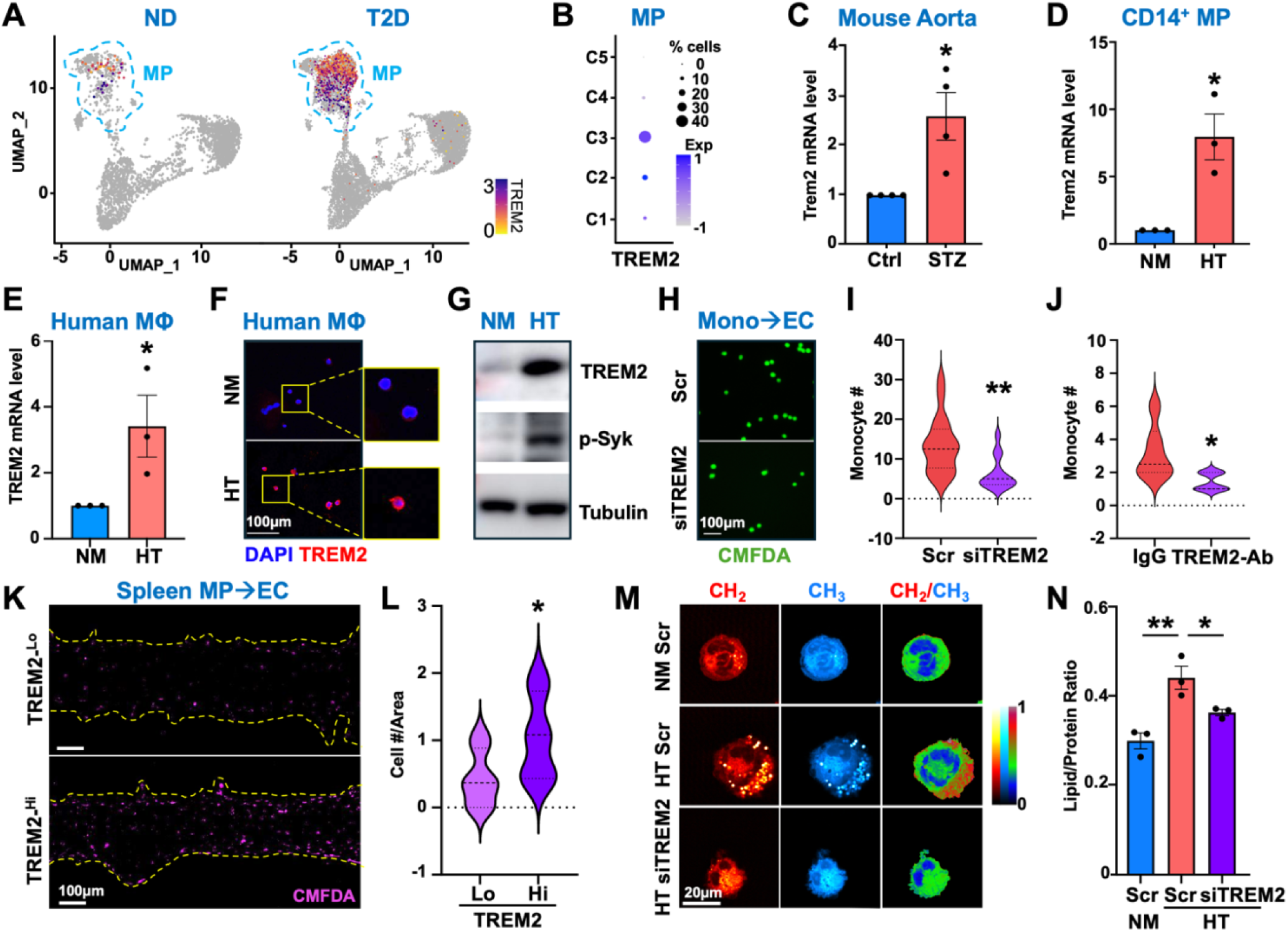
TREM2 is increased in DM-MP and promotes inflammatory response. (A,B) TREM2 expression in ND and T2D donors (in A) and in 5 MP clusters (C1-C5) (in B) quantified by intima scRNA-seq. **(C-E)** TREM2 mRNA levels in the aorta from male C57Bl/6 mice treated with STZ or citrate control (Ctrl) (n=4/group) (in C), human CD14^+^ MPs (in D), and THP-1-derived MΦ (in E) treated with 25 mM glucose plus TNF-α (HT) or 25 mM mannitol as osmolarity control (NM). **(F,G)** Immunofluorescence staining of TREM2 (with DAPI counterstain) and immunoblotting of TREM2 and p-Syk in THP1-derived MΦ as in (E). **(H-J)** Adhesion assay with ECs cultured under HT and incubated with CMFDA-labeled THP-1 cells with or without TREM2 knockdown (KD) (in H, I) or neutralizing antibody against TREM2, with IgG as isotype control (in J). **(K,L)** Adhesion assay with CMFDA labeled TREM2-high/low (TREM2-^Hi/Lo^) myeloid cells from two diabetic human spleens and HUVECs in a microfluidics chamber. Representative images (in K) and quantification of attached cells (in L). **(M,N)** THP-1-derived MΦs were treated as indicated. **(M)** Representative images with the first and second columns showing single-frequency label-free SRS targeted at lipid (–CH_2_) and the protein (–CH_3_) channels. The third column represents the corresponding ratiometric images of CH_2_/CH_3_ (lipid/protein). (**N**) Quantification of lipid:protein ratio. Data represent mean±SEM from 3-4 independent experiments and *p<0.05, **p<0.01 based on Student’s t-test (in C-E, I, J, L) or one-way ANOVA followed by Dunnett’s test in (N).

To determine the role of TREM2 in MP function under DM, we inhibited *TREM2* using siRNA or a neutralizing antibody (TREM2-Ab) (60, 61) in THP-1 monocytes. As a result of *TREM2* loss-of-function, either by knockdown (KD) at the RNA level (evident by decreased mRNA and protein levels) or block at the protein level (evident by decreased phospho-Syk) (**fig. S17**), the adhesion of monocytes to HT- exposed ECs was substantially decreased (**Fig. 4H-J**). In a complementary experiment, a TREM2 agonist (62) significantly increased monocyte adhesion to ECs in NM condition (**fig. S18**). In line with these findings, TREM2-high myeloid cells isolated from the spleen of human donors with DM (**fig. S19**) adhered significantly more to ECs in a microfluidic vessel-on-chip device (63), as compared to the TREM2-low counterparts (**Fig. 4K, L**).

TREM2 signaling is a critical to lipid metabolism in myeloid cells (64) and regulates lipid uptake in foamy MΦs in atherosclerosis (65). To examine the role of TREM2 in lipid uptake in the context of DM, we used stimulated Raman scattering (SRS) microscopy, which enables visualization of metabolites using a label-free imaging approach (66). SRS microscopy revealed an increased lipid-to-protein ratio in MΦ in HT as compared to NM, which was attenuated by TREM2 KD (**Fig. 4M, N**). Collectively, results in **Fig. 4** suggest that TREM2 plays a crucial role in the pro-inflammatory response and lipid uptake of MPs under diabetic conditions.

### TREM2 modulates EC function in diabetic conditions

Among all MP clusters identified from the intima scRNA-seq data from human mesenteric arteries, TREM2 is most abundant in MP-C3 (**Fig. 4B**), the cluster enriched for EC-relevant pathways, including angiogenesis, hypoxia response and nitric oxide biosynthetic process (**fig. S9**). Thus, we queried whether TREM2 expressed in MΦ can reciprocally regulate EC function under DM conditions. We treated human microvascular ECs (HMVECs) with conditioned medium (CM) from THP-1-derived MΦs with or without TREM2 KD and performed wound healing scratch assays (**Fig. 5A**). Compared to CM from NM-treated MФs, CM from HT-treated MФs reduced the wound closure of ECs. This inhibitory effect of HT-treated MФ CM was largely restored by TREM2 KD, with faster migration across most of the timepoints measured (**Fig. 5B, C**). Furthermore, CM from MФ under HT caused ECs to express higher levels of pro-inflammatory markers *VCAM1* and *ICAM1*, which was abolished with TREM2 KD in MФs (**Fig. 5D**). Of note, addition of HT alone to the media did not affect EC wound closure (**fig. S20**), suggesting that it is the secreted factors from HT-treated MФs, rather than HT per se, that impaired EC migration. Indeed, compared to the scrambled control, TREM2 KD decreased several pro-inflammatory cytokines, including IL-6, 7 and 8 (**Fig. 5E-G**). Collectively, these data suggest that TREM2-mediated signaling from MPs promotes inflammatory responses and inhibits wound healing capacities of ECs.

**Figure 5.**
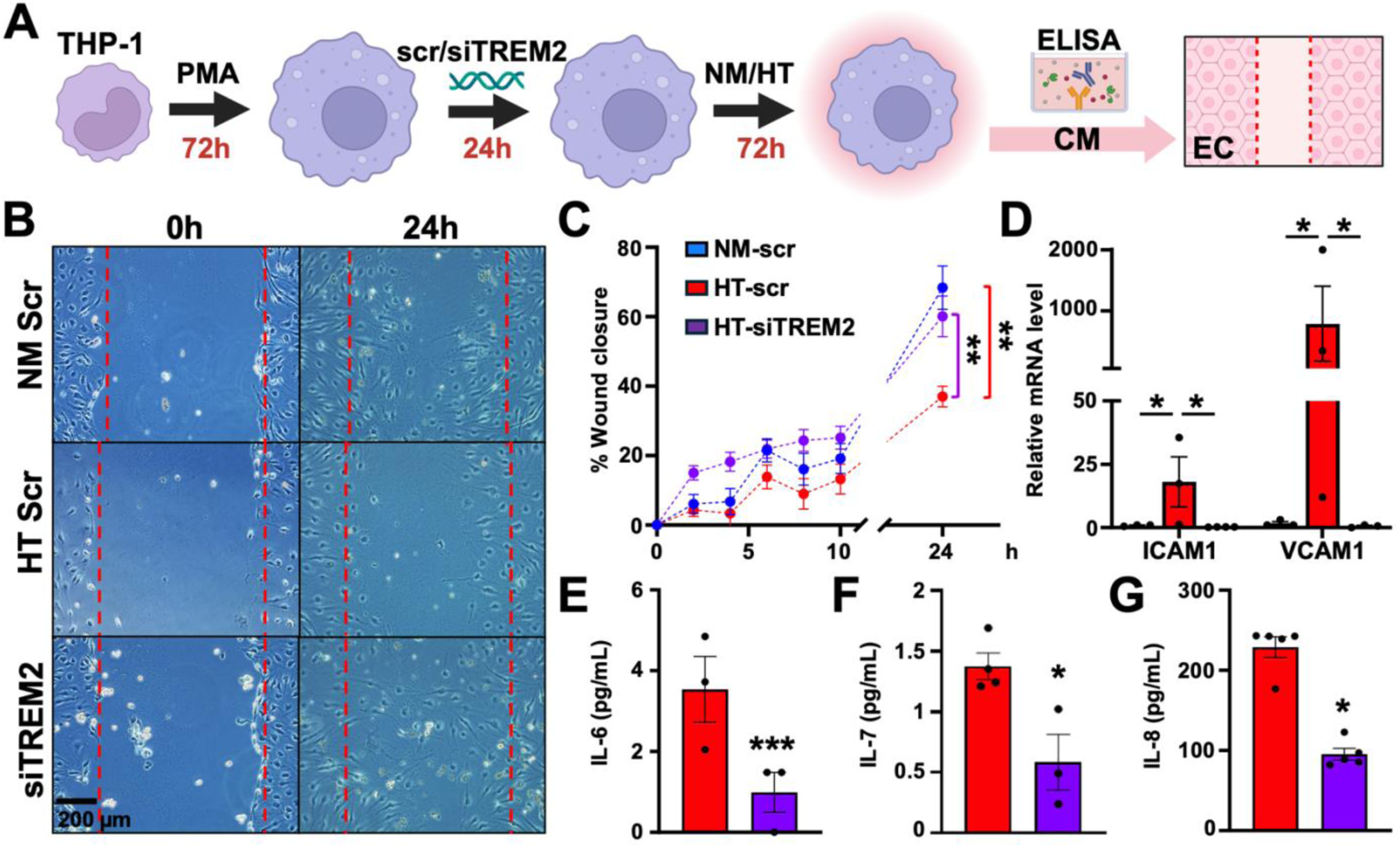
TREM2 modulates EC function in diabetic conditions. **(A)** Schematic of scratch wound assay using HMVECs and conditioned medium (CM) from THP-1-derived MΦ with or without TREM2 knockdown and treated by NM/HT. **(B,C)** Representative images and quantification of scratch wound assay using ECs incubated with MΦ CM across a 24 hour (h) time course. **(D)** qPCR of *VCAM1* and *ICAM1* in ECs treated with MΦ CM. **(E-G)** Levels of IL-6, 7, and 8 in MΦ CM as measured by Luminex. Data represents mean±SEM from 3-5 independent experiments. *p<0.05, **p<0.01, ***p<0.001 based on ANOVA followed by Dunnett’s post-hoc test (C, D) and Student’s t-test (E-G).

### TREM2 neutralization improves ischemic perfusion recovery in DM

Given the demonstrated role of TREM2 in pro-inflammatory responses and wound healing *in vitro*, we next investigated the role of TREM2 *in vivo* using a mouse model with DM and ischemic injury. Specifically, we induced hyperglycemia using STZ (which increases *Trem2* in the aorta [**Fig. 4C**]), followed by femoral artery ligation to induce HLI (**Fig. 6A, fig. S21A**). scRNA-seq of the limb muscles revealed that *Trem2* expression was increased in MФ in DM as compared to ND mice, and further increased in the ligated ischemic limbs as compared to the non-ischemic controls (**Fig. 6A**). In the diabetic mice, there was an increase in TREM2 protein and its co-localization with isolectin B4 ([IB4], EC marker) in the ischemic vs. non-ischemic limb muscles (**Fig. 6B, fig. S21B**). We then treated these mice with a local intramuscular (i.m.) injection of TREM2-Ab (same as used *in vitro*) (**Fig. 6C**). Compared to the IgG control, diabetic mice receiving TREM2-Ab showed enhanced recovery of perfusion (**Fig. 6D, E**), coupled with an increase in vascular density in the ischemic muscle (**Fig. 6F, fig. S22A**). Of note, in ND mice, TREM2-Ab did not significantly affect the recovery rate (**fig. S22B**). These data indicate that TREM2 is increased by ischemic injury under DM condition and TREM2 inhibition promotes angiogenesis and ischemic recovery.

**Figure 6.**
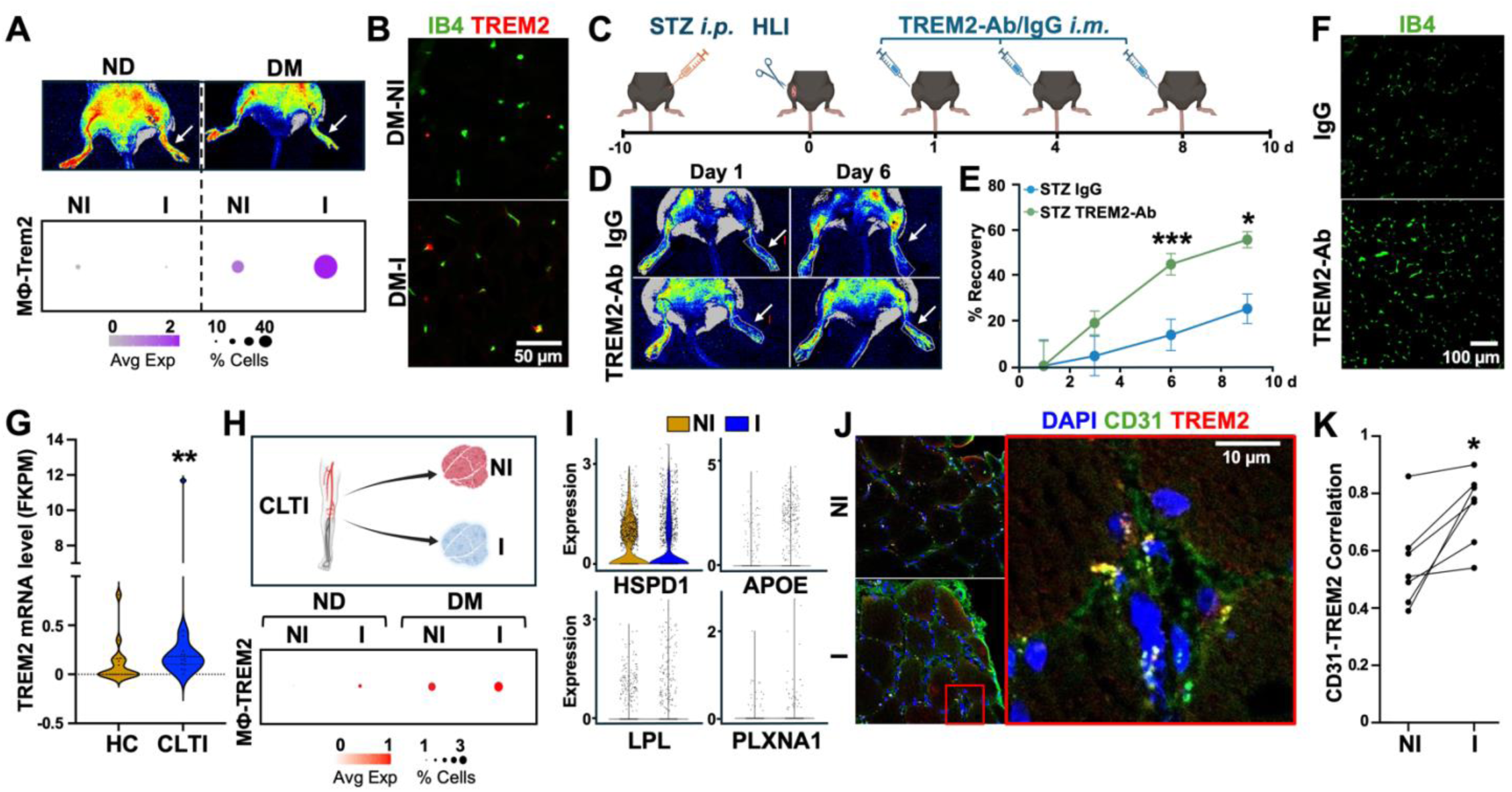
Induction of TREM2 in DM-PAD and its effect on ischemic recovery. **(A)** Top: Laser speckle images of flow perfusion in C57Bl/6 mice treated with or without STZ (DM vs ND), 5 days post hindlimb ischemia (HLI). Note the decreased flow in the ligated limb (white arrow) of DM mouse as compared to ND. Bottom: MΦ TREM2 expression in non-ligated (NI) or ligated (I) limb muscles from ND and DM mice, profiled by scRNA-seq (n=2/group). **(B)** IB4-TREM2 co-IF on the non-ligated vs ligated limb muscles from DM mice. **(C-F)** DM was induced in mice as in (A), followed by HLI and intramuscular injection of TREM2-Ab or IgG control on post-op Day (d) 1, 4, & 8. **(D,E)** Representative images of flow perfusion (in D) and % recovery quantified by comparing the flow perfusion in the ligated vs non-ligated limb of the same animals (in E). Data represented as mean±SEM of n=8/group (including 7 male and 9 female). **(F)** Representative images of IB4 staining of ischemic muscles on post-op Day 6. **(G)** TREM2 mRNA levels determined by bulk RNA-seq of human muscle biopsy from healthy control (HC, n=61) and patients with critical limb ischemia (CTLI, n=64). **(H,I)** scRNA-seq profiled expression of *TREM2* (in H) and TREM2 ligands in ECs (in I) in (non-)ischemic (NI/I) muscles from CLTI patients. **(J,K)** CD31-TREM2 co-IF in NI/I muscles from human patients with CLTI (n=7). Representative images (in J) and quantification (in H) of CD31-TREM2 co-staining expressed by Pearson’s R values. *p<0.05, **p<0.01 ***p<0.001 based on Student’s t-test.

### TREM2-EC interactions in human DM-PAD

To explore the disease relevance of TREM2 and its interactions with ECs in human PAD, which is partially modeled by the HLI in mice, we leveraged two published PAD RNA-seq datasets. One is bulk RNA-seq of gastrocnemius muscle from 32 healthy controls (HC) and 19 patients with CLTI (67), and the other scRNA-seq of (non-)ischemic muscles from three patients with CLTI (two ND and one T2D) undergoing endovascular procedures (68). *TREM2* was increased in 1) skeletal muscle from patients with CLTI vs HC (**Fig. 6G**) and 2) skeletal muscle from DM vs ND patients (**Fig. 6H**). Specifically, *TREM2* levels in the muscle from patients with CLTI follow this order: DM-ischemic > DM-non-ischemic > ND-ischemic > ND-non-ischemic (**Fig. 6H**), similar to the mouse DM-HLI model (**Fig. 6A**). In line with the increased levels of *TREM2* in ischemic vs non-ischemic muscles, those of TREM2 ligands in ECs were also higher in the ischemic muscle (**Fig. 6I**). Furthermore, co-staining of TREM2 and CD31 in muscle samples from 8 patients with CLTI (**Table S4**) showed more proximity in the ischemic sites, as compared to the non-ischemic sites, evident by the higher Pearson’s R value (**Fig. 6J, K**), suggesting more TREM2-EC interactions in ischemia. Taken together, these data demonstrate that TREM2 and its interaction with ECs are present and increased in human DM-PAD.

## Discussion

In this study, we leveraged scRNA- and ST-seq to investigate EC-MФ interactions within DM vasculopathies. By mapping DM-associated transcriptomic changes in human arteries at the single cell level within their native spatial context, we inferred key interactions between ECs and MPs (**Fig. 1-3**). A standout discovery was TREM2 **−** identified as the top DM-induced MФ receptor **−** which was used as an example to demonstrate how DM-induced MФ receptors engage in crosstalk with ECs. Through integrative analyses combining cell culture, a mouse model of DM-PAD, and samples from human donors with DM and PAD, TREM2 was identified as a critical mediator of EC-MФ communication (**Fig. 4-6**). Our study thus provides a valuable, high-resolution resource for dissecting DM-associated vascular pathology, highlighting a pivotal previously unknown role of TREM2-mediated EC-MФ communication in diabetic vasculopathy, with direct translational importance validated through findings in human PAD samples.

EC dysfunction and EC-MP interplay are key early events driving vascular inflammation and dysfunction. Consistent with this, human intima scRNA-seq revealed remarkable transcriptomic changes in ECs associated with DM (**Fig. 2A-C**). This was accompanied by an increase in intimal MPs (**Fig. 1E-G**), indicating activation of inflammatory response and enhanced EC-MP interactions in DM vessels (**Fig. 3C, fig. S15**). DM-associated changes in MPs were evident not only in DEGs but also in notable shifts in MP clusters (**Fig. 2F, G**). Among the five MP clusters identified, MP-C3 showed the greatest increase in T2D samples with the highest TREM2 expression (**Fig. 2G**, **4B**). This TREM2 increase in the vasculature with DM, previously unreported, was validated in cultured mouse and human MPs treated with HT, a hyperglycemia mouse model, and muscle samples from PAD patients with DM. Interestingly, TREM2 is also elevated in PAD patients (compared to HC) and ischemic muscles (compared to non-ischemic regions) in mouse and human (**Fig. 6**), suggesting TREM2 is a key molecule linking diabetes, vascular inflammation and ischemic disease such as PAD.

In line with the increased TREM2 expression in MPs in DM, we identified elevated TREM2-EC interactions within the diabetic vasculature. This finding is supported by scRNA-seq, ST-seq, and CD31-TREM2 co-staining of human arteries. To date, several putative TREM2 ligands across different tissues have been identified and the repertoire continues to grow (69). While the exact TREM2 ligands on ECs remain to be explored, several candidate ligands show increased pairing with TREM2 in T2D arteries (**Fig. 3D, fig. S15**). Given that TREM2 expression also rises during monocyte differentiation into MΦ (65) and our findings that TREM2 activation enhances monocyte adhesion to ECs under NM while TREM2 inhibition reduces monocyte adhesion under HT (**fig. S18, Fig. 4H-J**), we speculate that dysfunctional ECs in the diabetic vascular wall recruit monocytes, upregulating TREM2 to adhere to ECs and facilitate diapedesis and differentiation into MΦ.

To further examine TREM2’s role in MP and EC function in DM, we found that TREM2 inhibition in MΦ reduces lipid uptake and inflammatory cytokine production (**Fig. 4M, N, Fig. 5E-G**). Conditioned media from these TREM2-inhibited MΦ suppresses adhesion molecule expression in ECs while enhancing EC migration and wound healing capacity (**Fig. 5**). Moreover, TREM2 inhibition improved ischemic flow recovery in diabetic mice (**Fig. 6C-F**). Together, these findings suggest that TREM2 mediates a multi-step, bidirectional communication between ECs and MPs in DM, propagating inflammation and contributing to sustained vascular damage.

First identified in microglia, TREM2 has demonstrated both disease-protective and promoting roles. In the brain, TREM2 can promote the clearance of cellular debris to preserve neurological function (70–72), or drive a senescent microglial population to contribute to Alzheimer’s disease (73). In metabolic regulation, both overexpression and KO of TREM2 have been shown to promote obesity and insulin resistance (48, 74, 75). In cancers, TREM2 is highly expressed on tumor-associated MΦs to promote tumor growth and immune evasion (76). In the vasculature, the role of TREM2 has been explored primarily in atherosclerosis (46, 49, 65, 77, 78). Deletion of TREM2 has been shown to increase necrotic core formation in early atherosclerosis (49, 65, 79) and attenuate plaque progression in advanced lesions (65). Some of these seemingly inconsistent findings may stem from using global vs MP-specific TREM2-KO mice (48, 74, 75, 80, 81), given that other cells than MP may also express TREM2 (77). On another hand, these studies also underscore a context-dependent role of TREM2 (47). In support of this notion, TREM2-Ab treatment improved ischemic recovery in diabetic mice but had no significant effect in ND mice (**Fig. 6** and **fig. S22B**), likely due to the relatively low expression of TREM2. Additionally, a recent report suggests a protective role for TREM2-high MΦ in diabetic nephropathy (50). Our study, which points to a disease-promoting role for TREM2 in DM-associated PAD, emphasizes highly context-dependent roles of TREM2 in disease progression, which remains to be fully elucidated.

TREM2 has emerged as a promising therapeutic target across various diseases. Both TREM2 agonists and antagonists are being developed for potential use in treating Alzheimer’s disease and cancer (76, 82), among other diseases. Our findings from human donors and patients with DM and PAD, provide a strong link between EC dysfunction, EC-TREM2 interaction, and diabetic vasculopathy. Using a preclinical model, we showed inhibition of TREM2 leads to improved functional outcome of diabetic limb ischemia. These findings hold significant translational relevance. On one hand, our study supports a rationale of inhibiting TREM2 to treat inflammatory ischemic diseases, such as DM-PAD. On the other, it implies the need for caution in the therapeutics aimed at enhancing TREM2 function, particularly in patients with existing DM and ischemic disease.

Our studies have several limitations. First, we initially profiled human mesenteric arteries, which may not fully capture the DM-associated changes across other vascular beds. Given the complexity of donor backgrounds, some observed changes in T2D arteries could be influenced by other risk factors, such as hypertension or hyperlipidemia. Additionally, our use of local intramuscular injection of TREM2-Ab may have affected ischemic recovery through actions of cells other than MPs. Future studies are needed to further evaluate TREM2 as a therapeutic target for DM-PAD. Despite these limitations, our study provides an atlas of diabetic human arteries with single-cell and spatial resolution, identifies a novel and critical role of TREM2-EC interactions in diabetic vasculopathy, and serves as proof-of-principle for targeting TREM2 as a potential therapeutic approach for DM-PAD.

## Methods

### Human Tissues

Studies on mesenteric arteries were conducted on deidentified specimens obtained from the Southern California Islet Cell Resource Center at City of Hope. The research consents for the use of postmortem human tissues were obtained from the donors’ next of kin and ethical approval for this study was granted by the Institutional Review Board of City of Hope (IRB #01046). The selection of donors was based upon the Integrated Islet Distribution Program criteria. T2D was identified based on diagnosis in the donors’ medical records as well as the percentage of glycated hemoglobin A1c (HbA1c) of 6.5% or higher. For scRNA-seq, the intimal cells were isolated from human mesenteric artery by scraping the innermost layer with TrypLE as previously described (25). For scRNA-seq on whole arterial tissue, cells were obtained using a TrypLE and Liberase based digestion. Briefly, tissue was washed three times in PBS, and dissociated in an enzyme cocktail (2 U/mL, Liberase [5401127001, Sigma-Aldrich] and 2 U/mL elastase [LS002279, Worthington] in Hank’s Balanced Salt Solution [HBSS]), minced with scissors and incubated at 37 °C for 1 hour, the cell suspension was strained and then pelleted by centrifugation at 500g for 5 min. The enzyme solution was then discarded, and cells were resuspended in fresh HBSS. For Visium, whole artery was rapidly frozen in pre-cooled isopentane and stored in −80°C prior to use. For IHC, whole artery was embedded in paraffin.

PBMCs and CD14^+^ monocytes were isolated as described (58) from the peripheral blood of donors in accordance with a protocol approved by the City of Hope Institutional Review Board recruited under (IRB #00123).

Skeletal muscle was obtained from CLTI patients undergoing lower-limb amputation in accordance with a research protocol approved by the Duke University Institutional Review Board (IRB Pro00065709). Paired samples from proximal and distal muscle bodies were harvested and embedded in OCT using liquid nitrogen. For above knee amputations, the proximal muscle specimen was obtained from the vastus medialis and the distal specimen from the tibialis anterior. For below knee amputations, proximal and distal specimens were both obtained from the tibialis anterior. Sections of 8 μm diameter were prepared using cryostat sectioning for staining.

### Animal Study Approval

Animal experiments were approved by the Institutional Animal Care and Use Committees at City of Hope (#18069 & #17010) and Institutional Biosafety Committee (#16023).

### Statistical Methods

Statistical analyses for data other than high-throughput sequencing were performed using GraphPad Prism with input from a PhD level biostatistician from the Biostatistics Core at City of Hope. All data are representative of at least three independent experiments unless specifically noted. The sample size was precalculated to satisfy power requirements (with >85% confidence) for animal experiments. To accomplish randomization for imaging analysis and animal experiments, immunofluorescence, immunohistochemistry and perfusion images were sorted by a secondary blinded investigator prior to analysis. For sequencing experiments, genes were considered differentially expressed if their adjusted p value (q value) was below 0.05 and above a logFC threshold of 0.25. Two-group comparisons were performed using either unpaired or paired parametric Student’s t-test based on the normality of the data (specified in figure legends) and multiple-group comparisons were performed using ANOVA followed by post hoc tests as appropriate (specified in figure legends). *P* values less than 0.05 were considered statistically significant.

### Data and materials availability

All data needed to evaluate the conclusions in the paper are present in the paper and/or the Supplementary Materials and all high-throughput sequencing data used in this study will be deposited on GEO upon acceptance of this manuscript.

### Additional Methods

Additional detailed methods are provided in the supplementary material.

## Acknowledgments

The authors thank Prof. Marc K. Halushka at Cleveland Clinic for histological consultation of vascular tissues and valuable discussions and critiques, Dr. Yan Li at Case Western University for consultation on the use of RePACT for scRNA-seq analysis and Omisha Sangani at University of California, Riverside, for technical assistance. Schematics in **Figs. 1**, **5** and **6** were made with BioRender.

## Author contributions

Conceptualization: NKM, ZBC

Methodology: NKM, YL, XT, MQ

Investigation: NKM, YL, XT, RC, AT, DY, SY, XL, MC, MR

Supervision: LW, EL, RN, KWS, ZBC

Writing-original draft: NKM, ZBC

Writing-review & editing: NKM, YL, RC, AT, SY, XL, MC, LW, JPC, RN, ZBC

