## Supplementary Material for "Mapping Endothelial-Macrophage Interactions in Diabetic Vasculature: Role of TREM2 in Vascular Inflammation and Ischemic Response"

### Supplementary Materials

Supplementary Materials and Methods

Supplementary Figures (Fig. S1-22)

Supplementary Tables (Table S1-5)

### Supplementary Methods

#### Single-cell (sc) RNA-seq data analysis

Fastq files were first analyzed using Cell Ranger (v6.1.1), the standard pipeline provided by 10X Genomics, which aligned the raw reads to human hg38 reference transcriptome. The aligned data was then processed using the R package Seurat (v3.2.3) following published guidelines (83) and as we previously described (24, 28). Initially, well-established filtering steps were performed to remove cells expressing less than 200 genes and with high mitochondrial read percentages (>20%). Next, the "sctransform" normalization strategy was used where the residuals of negative binomial regression were used to model each gene. Datasets from the ten (for intima scRNA-seq) and three (for whole vessel scRNA-seq) samples then underwent integration to enable joint dimensionality reduction and clustering without being influenced by donor condition. Cells-pairs in matched biological states between samples were identified as "anchors" and used to integrate the data. The standard pre-processing step of scaling was performed prior to principal component analysis (PCA) on all cells and top 1,000 highly variable genes using the scaled z score expression values were identified. The top 15 significant principal components (PCs) were used as an input to the uniform manifold approximation and projection (UMAP) algorithm with 0.5 resolution (Seurat default). Cell clusters were generated and annotated using the following markers based on existing literature and PanglaoDB (34, 36, 41): for VSMCs, *ACTA2*, *MYH11*, *WT1*, and *TAGLN*; for ECs, *VWF*, *PECAM1*, *EGFL7*, *ID3*, *GNG11*, *MCAM*, *CLDN5*, and *CDH5*; for MP, *CD68*, *MARCH1*, and *CD52*; for fibroblasts, *LUM* and *DCN*; for T cells, *CD3D* and *CD3E*; and for NK cells, *KLRD1* and *NKG7*. Differential expression analysis was performed on UMI counts that had been normalized using "LogNormalize" and scaled. The Seurat default, non-parametric Wilcoxon test was used to analyze genes expressed in at least 10% of cells. The log foldchange (FC) of the average expression was thresholded at 0.25 and a pseudocount of 1 was added to the averaged expression values for differential gene expression analyses.

#### Monocle 3 trajectory analysis

Single-cell transcriptomic pseudotime analyses were performed using Monocle (v3 1.3.1) (84). Given that gene expression within this subset was normalized, the SCTransform-normalized expression matrix and corresponding metadata were extracted from the corresponding Seurat object. Metadata and SCT counts were used to create a cell_data_set object. To preserve clustering structure from previous analyses, we also extracted PCA/UMAP embeddings, cluster IDs, and cell type annotations from the processed Seurat object and inserted those into the corresponding slots of the cell_data_set object. Trajectories were then inferred using the learn_graph and order_cells functions using get_earliest_principal_node to define the root of the trajectory. Cells were then recolored on a UMAP as a reflection of pseudotime.

#### CellChat analysis

Cell communication analyses were carried out using the CellChat R package (v1.5.0) (85). We selected the CellChat human database (Interactions considered include secreted signaling, ECM-receptor, and cell-cell contacts). First, we extracted SCTransform-normalized counts from the integrated Seurat object. We created a Cellchat object for matrices from each disease status using the createCellChat() function. We subsequently identified overexpressed genes in each condition using the identifyOverExpressedInteractions. Communication probabilities were estimated with computeCommunProb and aggregated cell communication networks calculated with the aggregateNet function. We then merged ND and T2D Cellchat objects using the mergeCellChat function. Differential cell communication, particularly between EC-MP was calculated and visualized using netVisual_diffInteraction, netVisual_heatmap, netVisual_embedding, netVisual_chord_gene functions. Significantly enriched pathways were denoted as those with $p < 0.05$.

**CellphoneDB analysis**

To enable a systematic analysis of cell-cell communication molecules, we applied cell communication analysis based on the CellPhoneDB, a public repository of ligands, receptors, and their interactions (55), as we have described (26). Membrane, secreted, and peripheral proteins of the cluster of different time points were annotated. Significant mean and Cell Communication significance (p-value $< 0.05$) was calculated based on the interaction and the normalized cell matrix achieved by Seurat Normalization.

**RePACT**

Seurat objects were subset for ECs followed by rescaling and normalization. Disease status (i.e. ND or T2D) annotations from donors were embedded into the metadata. The tool scRNA.RePACT was used to generate pseudoindices of disease state for cells. Visualization was performed using all RePACT generated tools (42).

**Visium spatial transcriptome mapping of mesenteric arteries**

ST-seq was conducted using the Visium spatial gene expression slide and Reagent Kit (PN-1000185 and PN-1000186, 10x Genomics) as we have described (25). Briefly, the mesenteric artery was sectioned at 10 µm thickness onto the gene expression slide, making sure not to cross the fiducial frame. We then strictly followed the user guide (Rev G) to Hemoxylin and eosin (H&E) stain tissue, extract RNA and optimize permeabilization of 37°C for 18 min. The cDNA amplification was performed on a touch thermal cycler (C1000, BioRad). Visium spatial libraries were constructed using Visium spatial Library construction kit (10x Genomics, PN-1000190). H&E staining of the corresponding sections was imaged using a Zeiss Observer II light microscope at 10x using the tilescan feature.

Fastq files from sequencing were aligned to images and human hg38 transcriptome using 10X Genomics pipeline SpaceRanger v1.2.0. To manually annotate regions, we used LoupeBrowser 5 to cluster via histological layers and exported the data as .csv files. Next, we used Seurat v3 to perform standard quality control steps such as filtering voxels with >20% mitochondrial reads and zero counts. We normalized using SCTransform as variance in molecular counts per voxel can be substantial for spatial datasets and embedded histology cluster data per slice. To project whole vessel scRNA-seq onto Visium, we applied the 'anchor'-based integration workflow same as for scRNA-seq, but here it enables the probabilistic transfer of annotations from a reference to a query set. Using projection with scRNA-seq data from whole vessel, we generated prediction scores for each voxel of each cluster. SpatialFeaturePlot was used for data visualization.

**Mouse models**

To induce hyperglycemia, streptozotocin (STZ, 50 mg/kg to male, 75 mg/kg to female Sigma-Aldrich, USA, S0130) was administered via intraperitoneal injection to 8 - 12 week-old C57Bl/6 mice after 6 - 8 hours of fasting on 5 consecutive days. Blood glucose was measured 7 days post the first STZ injection, and mice with glucose levels ≥ 16.5 mmol/L (300 mg/dL) were considered diabetic and used for subsequent experiments.

HLI was induced as described previously (86) with modifications. The surgical site was shaved and treated with topical antiseptic. The left femoral artery was isolated and occluded 5–6 mm distal to the inguinal ligament by ligation with 6-0 surgical silk. The control limb on the right side underwent a sham surgery without arterial ligation. Blood perfusion was measured by laser speckle flowgraphy using a PERICAM PSI Z system (Perimed AB). The ratio of blood flow in the ischemic to non-ischemic hindlimb was calculated and expressed as the perfusion recovery rate. Intramuscular injections of 0.3 mg/kg TREM2-Ab (MAB17291, R&D systems) or rat IgG isotype control (MAB0061, R&D systems) were administered at stated timepoints.

For scRNA-seq of the muscles, tissues were digested with DMEM containing Type I collagenase and ECs were enriched (1 mg/mL) by anti-CD144-conjugated magnetic beads and MACS columns (130-042-401, Miltenyi Biotec), following a protocol we have previously described (26).

#### Immunohistochemistry (IHC) and immunofluorescence (IF)

Histological examination of human mesenteric arteries was primarily processed by the Solid Tumor Pathology Core at City of Hope using antibodies against CD68 (790-2931, Ventana), CD31 (AC-0083A, Eptomics) and TREM2 (91068, Cell Signaling Technology). CD68-Yellow/CD31-Purple/TREM2-Teal triplex IHC staining was performed on Ventana Discovery Ultra IHC Auto stainer (Ventana Medical Systems, Roche Diagnostics, Indianapolis, USA). Briefly, the slides were loaded on the machine, and deparaffinization, rehydration, endogenous peroxidase activity inhibition and antigen retrieval were performed. The three antigens were sequentially detected, and heat inactivation was performed to prevent any cross-reactivity between the three antigens. Following each primary antibody incubation, DISCOVERY anti-Rabbit NP and DISCOVERY anti-NP-AP or DISCOVERY anti-Rabbit HQ and DISCOVERY anti-HQ-HRP were incubated. The stains were visualized by DISCOVERY Yellow Kit, DISCOVERY Purple Kit, and DISCOVERY Teal Kit (Ventana), respectively; counterstained with hematoxylin (Ventana) and coverslipped. Whole slide images were acquired with NanoZoomer S360 Digital Slide Scanner (Hamamatsu).

Skeletal muscle from mice was embedded in OCT and sectioned via a cryostat. Slides were fixed in 4% (vol/vol) paraformaldehyde (PFA) for 10 min, and permeabilized in 0.2% (vol/vol) Triton X-100 for 5 min. A blocking buffer [0.22-mm filtered 1% (wt/vol) BSA 0.1% (vol/vol) Triton X-100 PBS] was used to block nonspecific binding for 2 hours at room temperature. For immunofluorescent (IF) staining, antibodies against CD31 (50245725, Fisher Scientific, 1:100 dilution); IB4 fluorescein (FL1201-.5, Vector laboratories, 1:250 dilution) and TREM2 (702886, ThermoFisher, 1:100) were used as primary antibodies. As secondary antibodies, Alexa Fluor 555-conjugated goat anti-rat IgG (A-11007, Invitrogen, 1:200 dilution) or Alexa Fluor 488-conjugated goat anti-rabbit IgG (A-11037, Invitrogen, 1:200 dilution) were used as appropriate. Nuclei were stained with DAPI (P36935, Invitrogen). Images were taken using a ZEISS Axio Observer or ZEISS LSM 700 confocal microscope. To measure vascular density in the skeletal muscles, the number of IB4-positive spots across the images were quantified on ImageJ in a blinded fashion. Tissue autofluorescence was used to visualize and count the muscle fibers. Vascularity was determined by taking the average of IB4-positive spots as a fraction of the number of muscle fibers for 3-5 images per mouse.

IF was also performed with cells cultured on coverslips (Bioptechs) following the same protocol as for the tissues. For colocalization analysis, Pearson's R value was generated by use of ImageJ Coloc2 plug-in a blinded fashion. In brief, background staining was subtracted from each channel of interest and the pixel intensity correlation was determined.

#### Cell culture, treatments, and assays

All cells were kept under standard cell culture conditions (humidified atmosphere, 5% $CO_2$, 37°C). THP-1 cells (American Type Culture Collection [ATCC]) were cultured in RPMI-1640 (11875093, Fisher) supplemented with 10% fetal bovine serum (FBS) (PI23209, Fisher]. PMA at 20 ng/mL was added to the culture medium to enable differentiation to MΦs. HT condition was created by adding D-glucose (D16-1, Fisher Scientific) to a final concentration of 25 mM and TNF-α (PHC3015, Thermo Fisher

Scientific) to a final concentration of 5 ng/mL. As normal glucose (NM)/osmolarity control, D-mannitol (M120-500, Fisher Scientific) was added at 20 mM in medium with 5 mM glucose. In some experiments, cells were transfected with a scrambled control or ON-TARGETplus SMARTpool TREM2 siRNA (54209, Dharmacon) at 100 nM using Lipofectamine-RNAiMAX transfection reagent (13778150, Thermo Fisher Scientific) in Opti-MEM (31985070, Fisher) according to manufacturer's protocol.

HUVECs tested negative for mycoplasma contamination (200-05n, Cell Applications) were cultured in complete M199 medium supplemented with 20% FBS (M2520, Sigma) and 1× antibiotics (penicillin–streptomycin, 15140122, Gibco,) and subjected to NM or HT. For the cell adhesion assay, The THP-1 cells were labeled with CellTracker Green CMFDA Dye (Thermo Fisher Scientific) and incubated with monolayer HUVECs ($4 \times 10^3$ cells per $cm^2$) for 20 min in a cell culture incubator. The nonattached monocytes were washed off and the attached monocytes were imaged using an Echo Revolve microscope using the 10x lens and green fluorescent channel. Average numbers per condition were calculated using a counter on ImageJ in a blinded fashion from five randomly selected fields of technical duplicates.

For the scratch wounding assay, human microvascular endothelial cells (HMVEC [Cell Applications, 100-5n) were seeded onto 12-well plates and grown to confluence in Human Microvascular Endothelial Cell Growth Medium (111-500; Cell Applications). Cell monolayers were wounded with a 1000-μL pipette tip to generate a cut of ~1 mm in width. After two washing steps, cells were incubated with conditioned media. Images were taken at various timepoints and area lacking cells were determined by measurement using ImageJ in a blinded fashion.

**Isolation and differentiation of human $CD14^+$ monocytes**

Human $CD14^+$ monocytes were isolated from the blood of healthy volunteers as described before (58). Briefly, PBMCs were purified from whole blood on Ficoll-gradients, and $CD14^+$ monocytes were isolated by immunomagnetic negative selection using EasySep Human Monocyte Isolation Kit (19059, Stemcell Technologies). Human primary monocytes were differentiated into MΦ using M-CSF1 (50 ng/mL) for up to 1 week. Cells were cultured in RPMI containing 10% FBS, penicillin/streptomycin (Pen/Strep; 100 U/100 µg/mL), 2 mM glutamine, and 5.5 mM glucose. Where indicated, cells were treated with NM or HT. Cells were then lysed in QIAzol (Qiagen) for RNA extraction.

**Isolation of $TREM2^{-Hi}$ and $TREM2^{-Lo}$ cells from human spleen**

In a tube with 1 mL ice cold PBS (70011069, Fisher), approximately 1g of human spleen was thoroughly minced on ice using autoclaved scissors. PBS was added to the homogenate before passing through a 70 µm strainer (22363548, Fisher) and centrifuged at 600xg for 5 min at 4 °C. The supernatant was discarded, and the pellet was resuspended in ACK lysis buffer (A1049201, Fisher) and incubated at room temperature for 5 min before centrifuging again at 600xg for 5 min at 4 °C. Following 2 washes with PBS containing 2% FBS, cells were passed through a 40 µm strainer, centrifuged and washed with PBS. Next, we proceeded with FACs using rat anti-TREM2 (MAB17291, R&D systems, 1:100 dilution) in PBS containing 2% BSA and a secondary antibody Alexa Fluor 488-conjugated goat anti-rabbit IgG (A-11037, Invitrogen, 1:200 dilution), with all incubations on a rotator for 30 min at room temperature. Nuclei were stained with DAPI (P36935, Invitrogen). Cells were sorted on The Aria II SORP (Becton Dickinson) to assess for myeloid populations and gating strategies.

**Microfluidics**

Microfluidic vessel-on-chip devices were fabricated using soft lithography as described (87). The device consists of a polydimethylsiloxane (PDMS) housing and two parallel cylindrical microchannels within the 3D collagen hydrogel. PDMS (Sylgard 184, Dow-Corning, DC2065622) was mixed with a curing agent, provided in the Sylgard 184 PDMS kit, at 10:1 weight ratio (base:curing agent) and cured overnight at 60 °C on a silicon master. The PDMS was removed from the silicon master, trimmed, and surface activated by plasma etching for PDMS bonding to a cover glass (22 × 40 mm, Fisher, 12-545-C). Four reservoir walls were embedded in the PDMS (not in collagen 1) by making four holes through the PDMS device using a biopsy punch (with a 6-mm diameter). Bonding of the PDMS to the glass was followed by curing at 80 °C overnight for permanent bonding. The devices were plasma etched, treated with 0.01% poly-L-lysine (Sigma, P8920) for 1 hour, followed by 1% glutaraldehyde (Electron Microscopy Sciences, 16310) for 30 min and then rinsed three times with sterile water, and incubated overnight in sterile water at room temperature. Steel acupuncture needles (0.25 × 50 mm, Hawato, HS 25 × 50) were sterilized with 70% ethanol and BSA-coated (1 mg/mL in PBS), then introduced into the devices as a scaffold of a "casting method." The needle-inserted devices were air-gun dried and UV sterilized for 30 min. Collagen type I (Corning, 356236) was buffered with PBS, titrated to a pH of 8.0 with NaOH (Sigma), yielding a final concentration of 2.5 mg/mL, that was pipetted into the microfluidic devices and polymerized for 50 min at 37 °C. Cell growth medium was then added to the devices overnight, and needles were carefully removed to create 250 µm diameter channels in the collagen gel. After overnight incubation with cell growth medium, the devices were seeded with HUVECs. Briefly, enzymatically dislodged (0.05% Trypsin/EDTA) HUVECs were resuspended at $1 \times 10^6$ cells/mL in M199 media, and 100 µL of cell suspension was introduced into the right-side channel of the device via the media reservoirs that are connected to the right-side channel, allowing cells to adhere to the collagen matrix for 10 min before washing with growth medium. The devices were incubated for 2 days on a rocking platform in a humidified tissue culture incubator (5% $CO_2$, 37 °C), replenishing culture media daily. Devices were treated with 25 mM D-glucose plus 5 ng/mL TNF-α for 24 hours. FACs isolated TREM2-$^{Hi}$ and TREM2-$^{Lo}$ spleen cells were stained with 1-2 µM CMFDA dye for 30 min before introducing them to the HUVEC lumens through the media reservoirs and allowed to adhere between 1-3 hours. Next, the devices were fixed with 4% paraformaldehyde (Electron Microscopy Sciences, 15710) for 1 hour at room temperature and stained with DAPI. Chambers were imaged using a Leica HC FLUOTAR L40 x/0.95 water-immersion objective on an inverted Leica SP8 confocal microscope, and spleen cell number over area was quantified using ImageJ analysis.

**Stimulated Raman Scattering (SRS) microscopy and data analysis**

The setup of the SRS microscopy system is the same as reported previously (88). Briefly, an 80 µs pixel dwell time was used to acquire images which gave an image acquisition speed of 8.52 s/frame for a 320x320 pixel field of view (FOV) at a resolution of 0.497 µm/pixel. The wavelength of the pump beam was set to 791.3 nm and 797.3 nm for $–CH_3$ (2940 $cm^{-1}$) and $–CH_2$ (2845 $cm^{-1}$) channels, respectively. Images were analyzed and color-coded using ImageJ. For $CH_2$:$CH_3$ imaging, a mask image was generated by adjusting the threshold followed by normalization of non-zero values to one. Subsequently, $CH_2$ images were divided by $CH_3$ images from the same cell set. The resulting ratiometric image was then multiplied with the mask image to create the final ratio ($CH_2$:$CH_3$) image. The contrast was set from 0 to 1. All experiments were repeated with at least three technical and three biological replicates.

**Western blotting**

Skeletal muscle or cells were lysed with protein extraction reagent (NP40 Cell Lysis Buffer; Thermo Fisher) in the presence of a protease inhibitor (8340; Sigma-Aldrich) and a phosphatase inhibitor cocktail (5870; Cell Signaling). Tissue extracts (30 µg) and a protein ladder were loaded on an 8% SDS-PAGE gel followed by transferring onto a PVDF membrane. The membranes were blocked with 5% skim milk or BSA and were incubated with primary antibodies at 1:1000 dilution in 3% BSA overnight at 4 °C. The primary antibodies used were rabbit anti-TREM2 (PA581933, Fisher), rabbit anti-TREM2 (A10482, AbClonal), rabbit anti-Phospho-Syk (C87C1, Cell Signaling) and rabbit anti-tubulin (2146S, Cell Signaling). The membranes were then incubated with anti-rabbit (7074S, Cell Signaling) or anti-mouse (7076S, Cell Signaling) or anti-rat (AB297057, Abcam) HRP-conjugated secondary antibody at room temperature for 1 hour and developed using ECL substrate (34580, ThermoFisher).

### RNA extraction and quantitative PCR

Total RNA was extracted from cells or tissues using TRIzol reagent (Thermo Fisher Scientific) with an additional bead homogenization step when processing tissues. cDNAs were synthesized using PrimeScript™ RT Master Mix containing both Oligo-dT primer and random hexamer primers (RR036B, TAKARA). qPCR was performed with iTaq™ Universal SYBR® Green Supermix (1725122, Bio-Rad) following the manufacturer's suggested protocol, using the Bio-Rad CFX Connect Real Time system. β-actin was used as the internal control in human and 36B4 in mouse samples. A complete list of primer sequences is included in **Table S5**.

### Luminex assay

A Luminex X-MAP bead array assay was utilized to analyze cytokine levels in conditioned medium. Conditioned medium samples were centrifuged to remove debris and incubated with antibody-coupled magnetic beads (using curated human 14-plex panel [NB1097; Bio-techne]). A standard curve using known concentrations of analytes was prepared for quantification. After incubation and washing, detection antibodies and streptavidin-PE were added. The samples were analyzed using a Luminex 200 system, a multiplexing flow-based analyzer equipped with dual lasers and optics for bead and fluorophore detection, to quantify fluorescence and determine analyte concentrations based on the standard curve.

**Supplementary Figures**

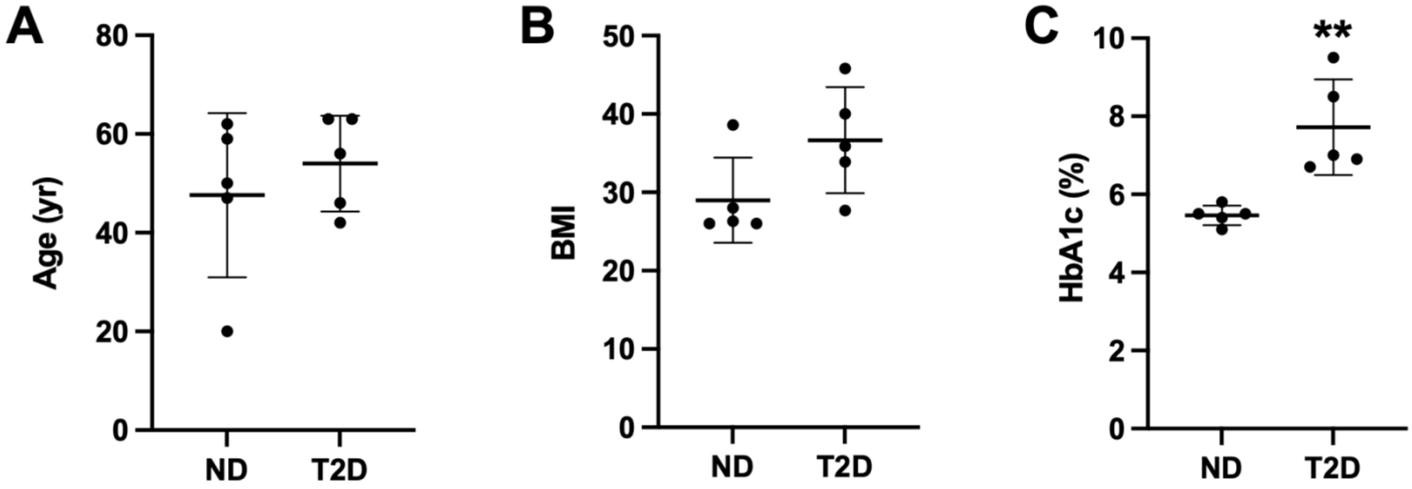


**Fig. S1. Demographics of human donors used for intima scRNA-seq.** Age (in A), BMI (in B), and HbA1c (%) levels (in C) of 5 ND and 5 T2D donors. Data represented as mean±SEM. **$p<0.01$ compared to ND based on Student's t-test.

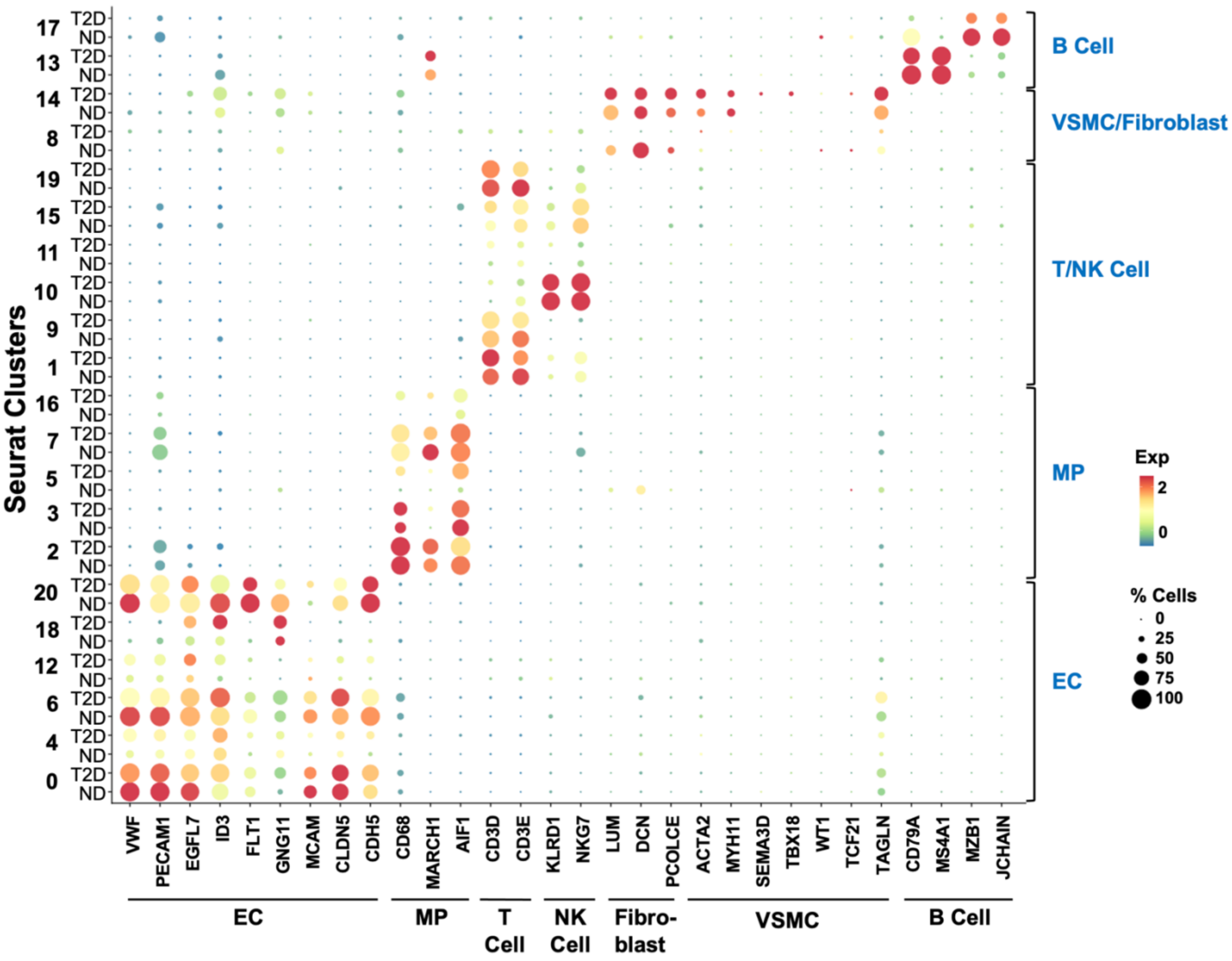


**Fig. S2. Expression of marker genes for cell annotation in 21 clusters.** Twenty-one clusters (Cluster 0-20) were identified from the intima scRNA-seq by unsupervised clustering through Seurat. Dotplot shows the percentage and average expression of cell marker genes (on x-axis) in each cluster split between ND and T2D (on y-axis).

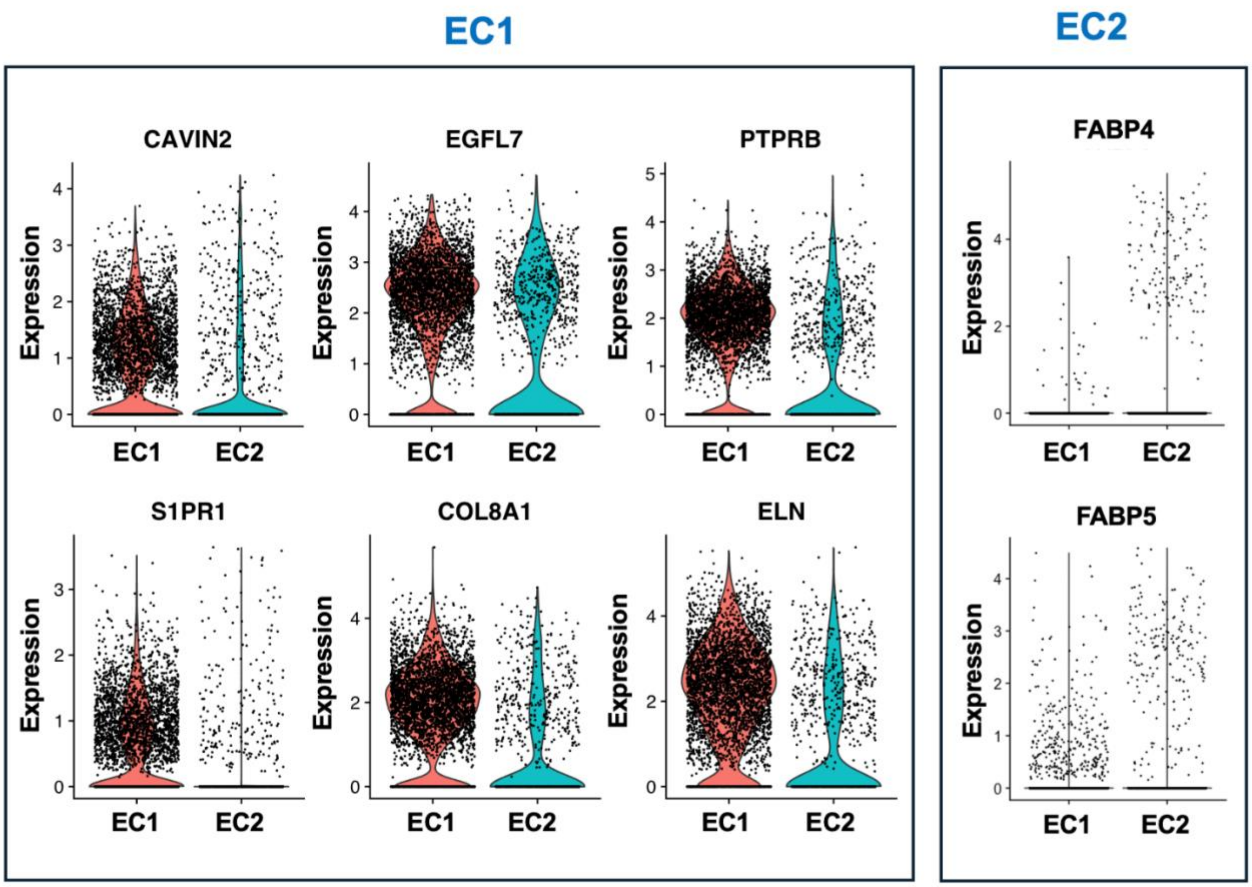


**Fig. S3. Characteristics of the two EC clusters.** Violin plots of gene expression as indicated in the two EC clusters, i.e., EC1 and EC2 quantified by scRNA-seq.

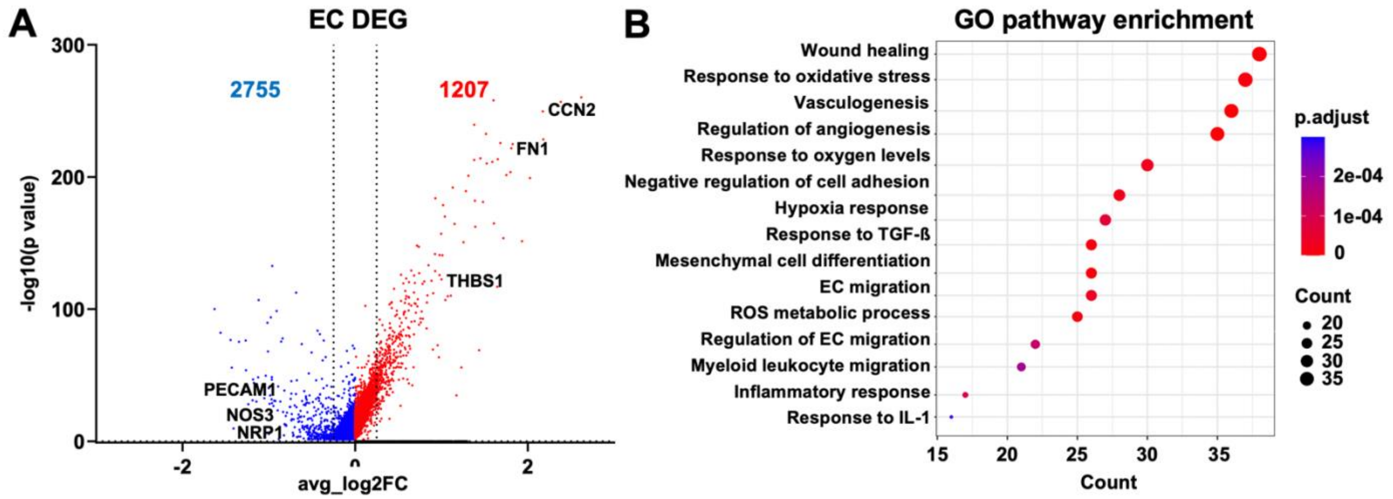


**Fig. S4. Single-cell transcriptomic changes in ND vs T2D ECs. (A)** Expression of DEGs between ND and T2D in ECs (blue: downregulated and red: upregulated) identified from intima scRNA-seq plotted with average log2 FC and -log10 p value. Dashed lines represent log2 FC cutoff (±0.25). **(B)** Dotplot of representative enriched pathways in EC-DEGs between ND and T2D, ranked by gene count.

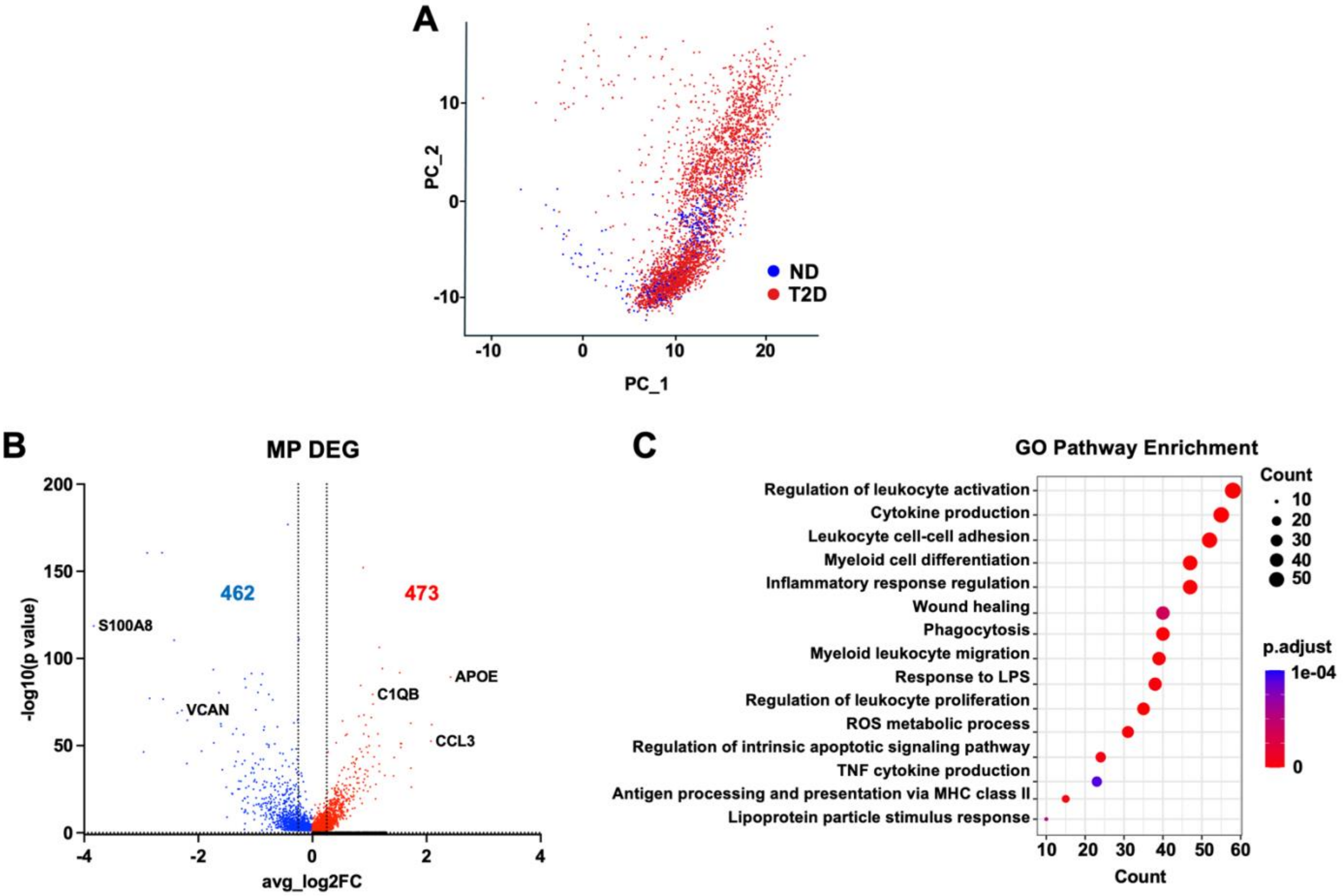


**Fig. S5. Single-cell transcriptomic changes in ND vs T2D MPs. (A)** PCA plot of MPs, separated by ND vs T2D generated by Seurat. **(B)** Expression of DEGs between ND and T2D in MPs (blue: downregulated and red: upregulated) identified from intima scRNA-seq plotted with average log2 FC and -log10 p value. Dashed lines represent log2FC cutoff (±0.25). **(C)** Dotplot of representative enriched pathways in MP-DEGs between ND and T2D, ranked by gene count.

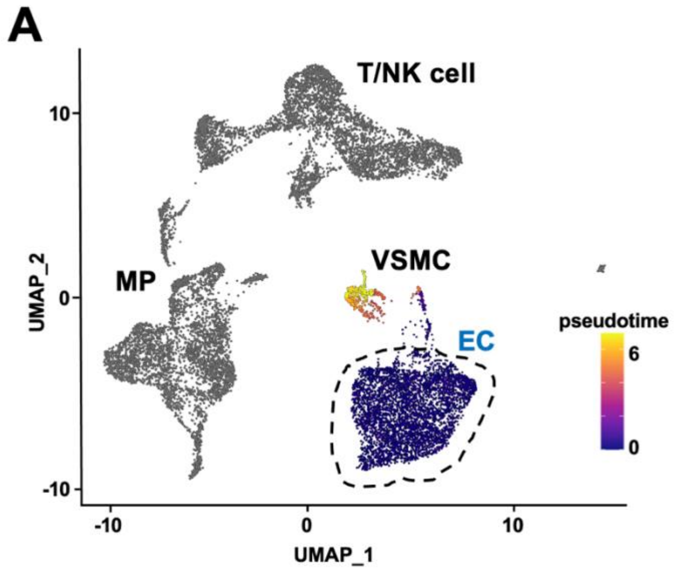


**Fig. S6. Trajectory analysis of intima scRNA-seq data.** Monocle 3 generated trajectories superimposed on UMAP of intima scRNA-seq, showing EC-rooted (in A) and T/NK cell-rooted trajectories (in B), with highlighted cells colored by pseudotime.

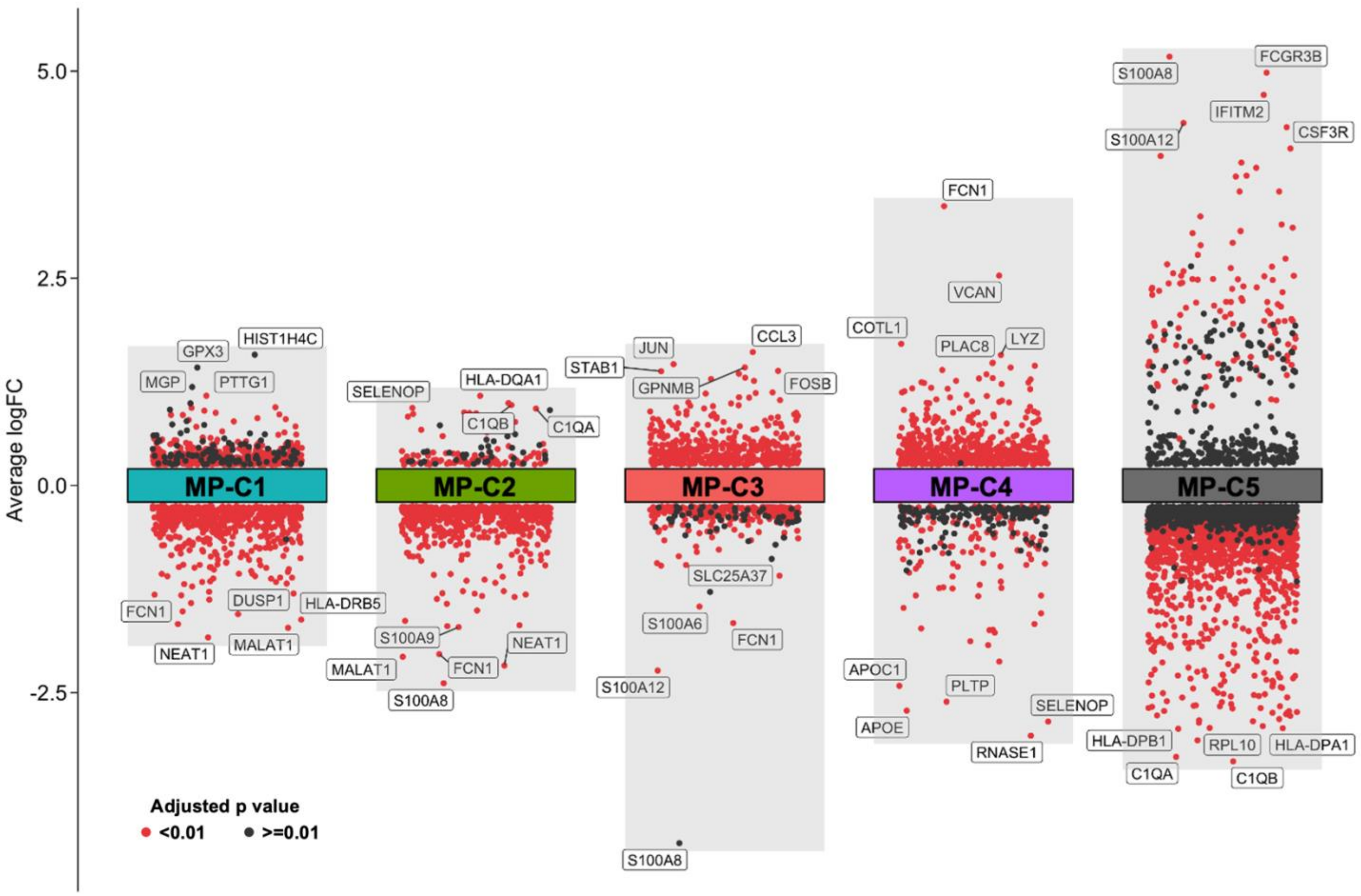


**Fig. S7. DEG features of MP subclusters.** Jitterplot showing DEGs among MP-C1 to C5 plotted with logFC of expression quantified by intima scRNA-seq. The color of the dots denotes p value. Red indicates significant DEGs and black indicates non-significant DEGs in given MP clusters.

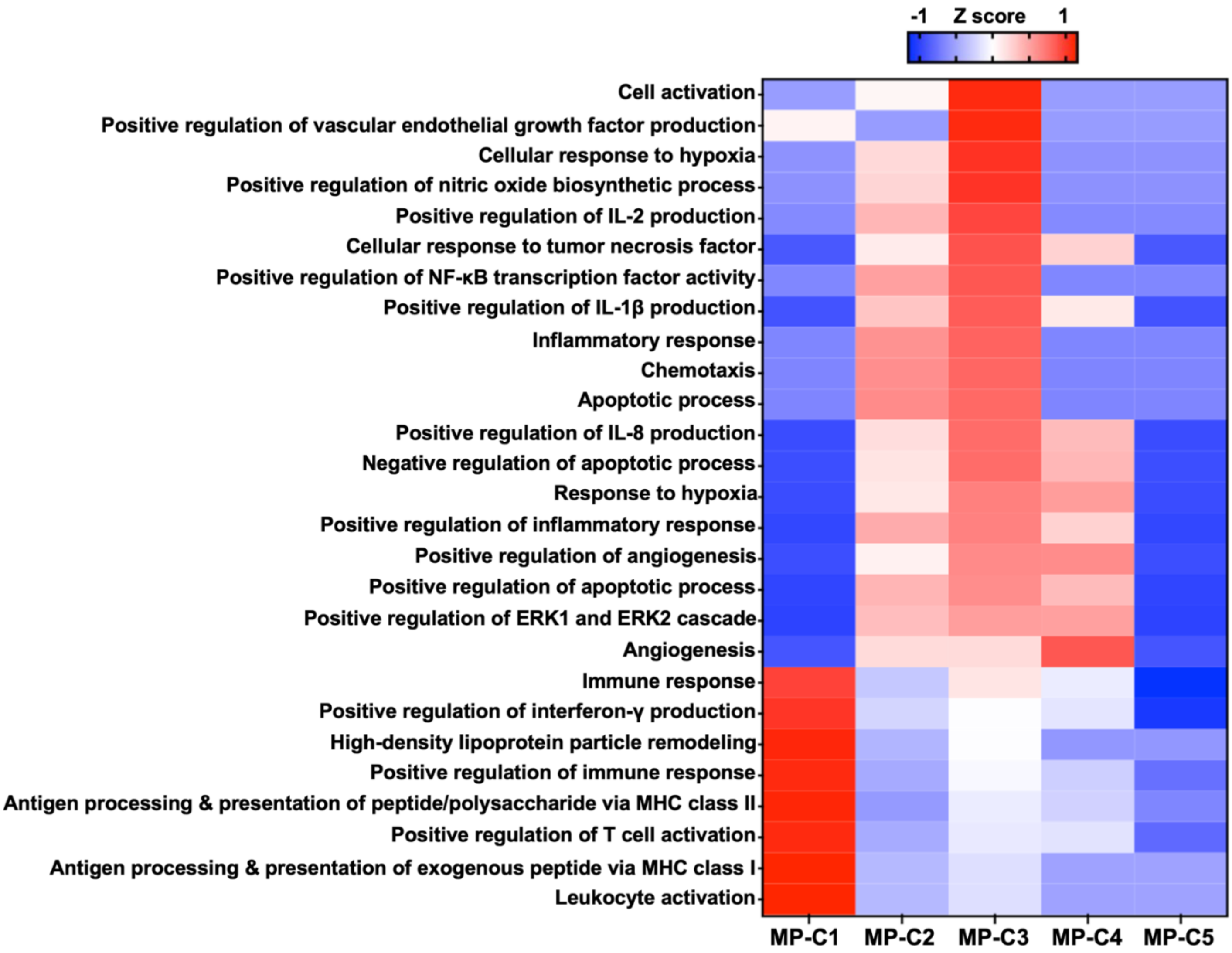


**Fig. S8. Enriched pathways in MP clusters.** Heatmap of enriched pathways between ND and T2D in MP-C1 to C5, plotted with Z scores of p values and ranked by the degree of enrichment in MP-C3.

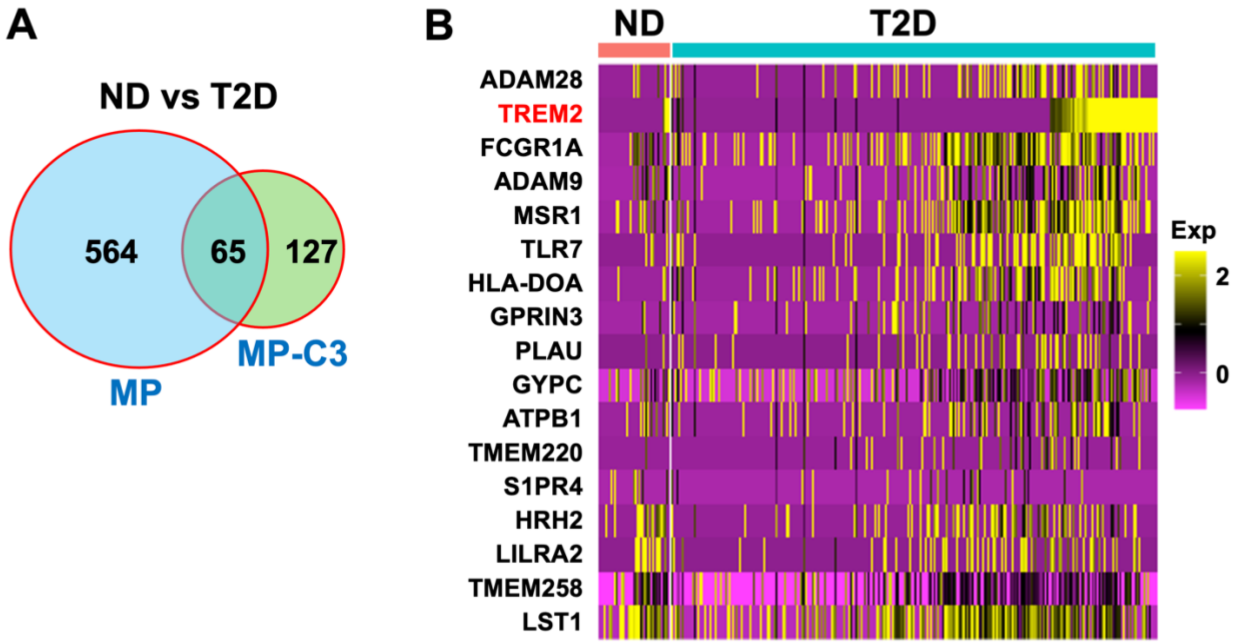


**Fig. S9. T2D-associated DEGs in MP-C3. (A)** Venn diagram of 629 DEGs in MP (vs all other cell types) between ND vs T2D (left) against 192 DEGs unique to MP-C3 (vs MP-C1, MP-C2, MP-C4, MP-C5), with 65 T2D-associated DEGs overlapping between all MPs and MP-C3. **(B)** List of top-ranked DEGs encoding cell surface proteins for both MP and MP-C3. Note that TREM2 is second-ranked by Surfaceome score and the top-ranked receptor.

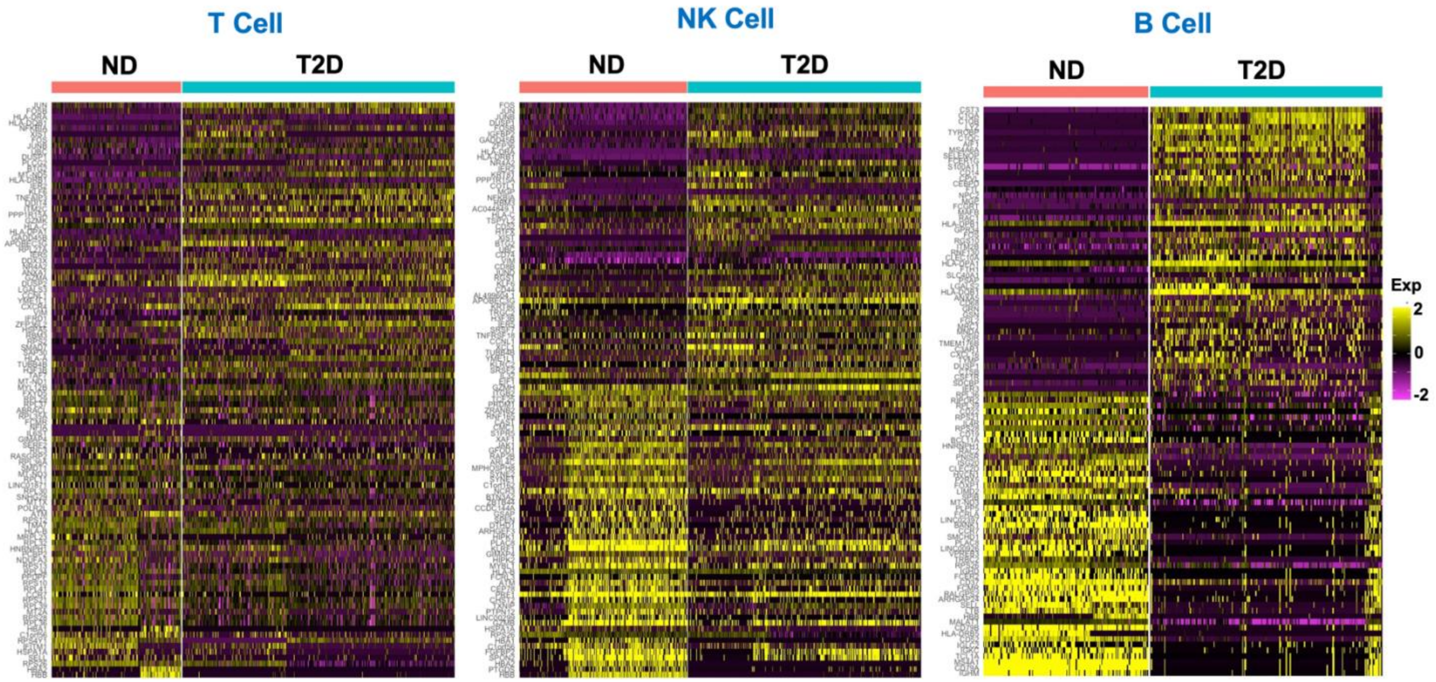


**Fig. S10. T2D-associated transcriptomic changes in other immune cell types.** Heatmaps of top 50 up- and down-regulated DEGs in T, NK, and B cells between ND and T2D revealed by intima scRNA-seq, plotted with Seurat and ranked by log2 FC.

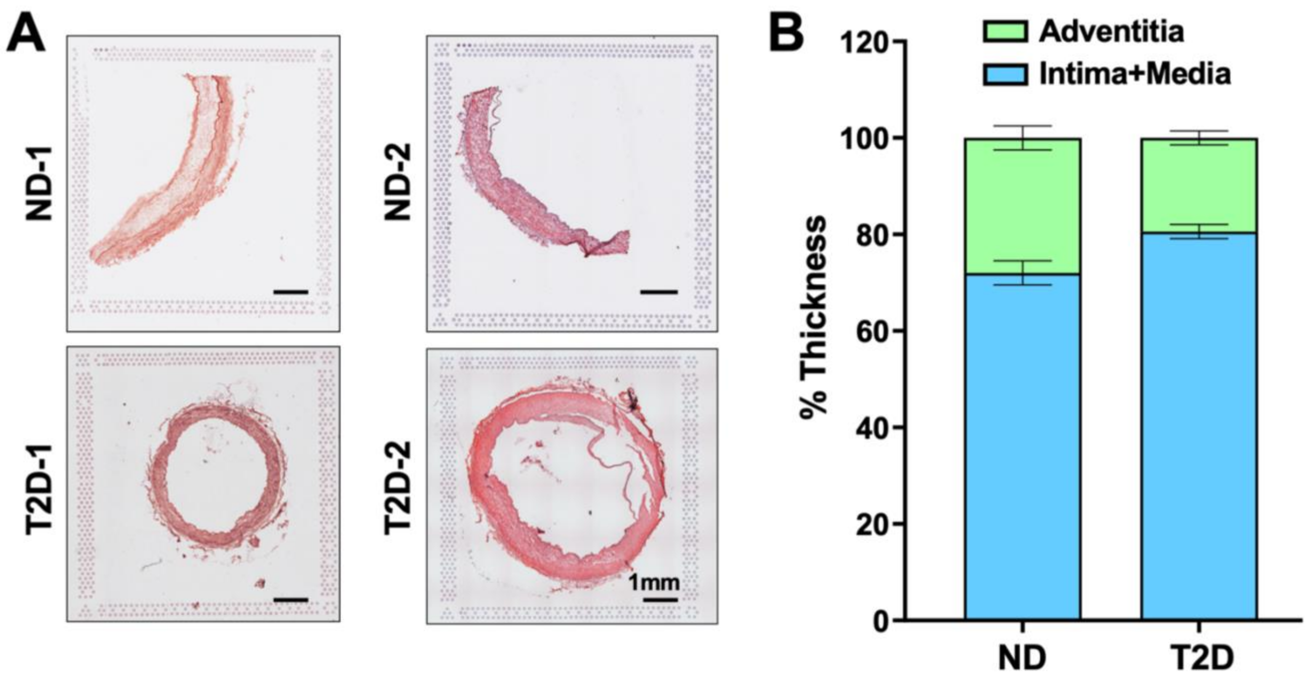


**Fig. S11. Histology of vessels for ST-seq. (A)** H&E staining of cross-sections and **(B)** quantification of intima-media thickness as a percentage of arterial wall thickness of 4 mesenteric arteries (2 ND and 2 T2D) used for Visium.

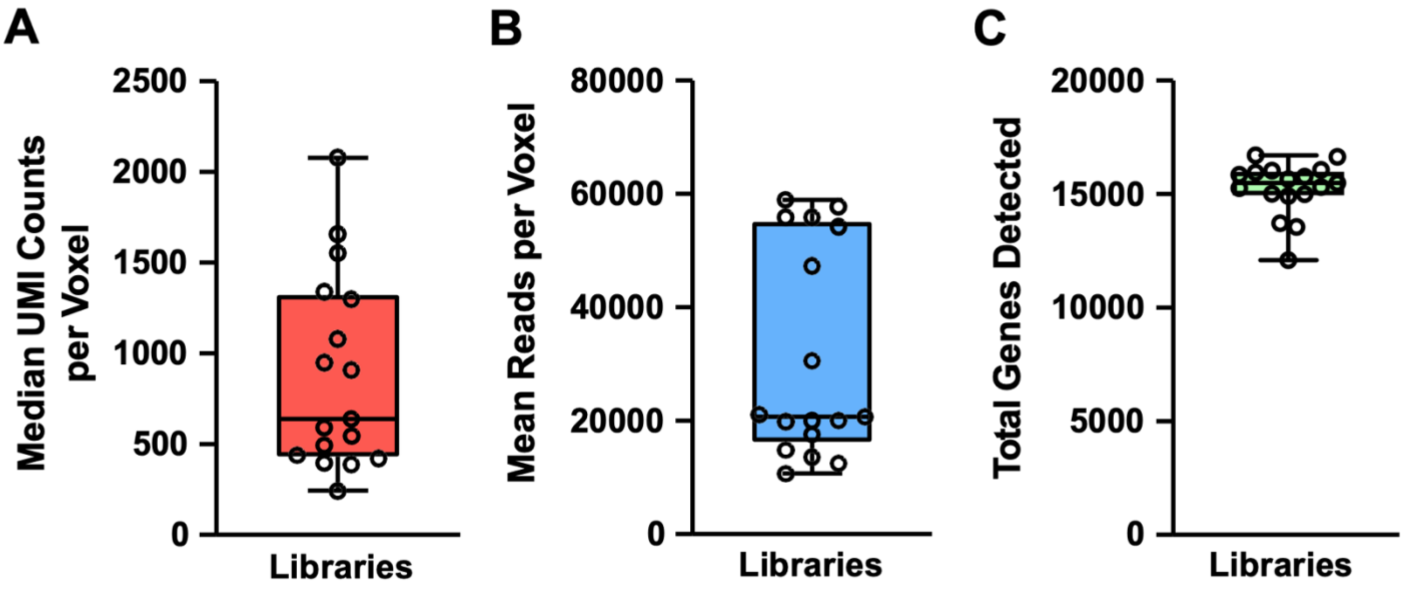


**Fig. S12. QC metrics of Visium experiment**. Output metrics from SpaceRanger show ranges between average **(A)** UMIs, **(B)** reads, and **(C)** genes per voxel for different samples/libraries. Error bars denote min and max.

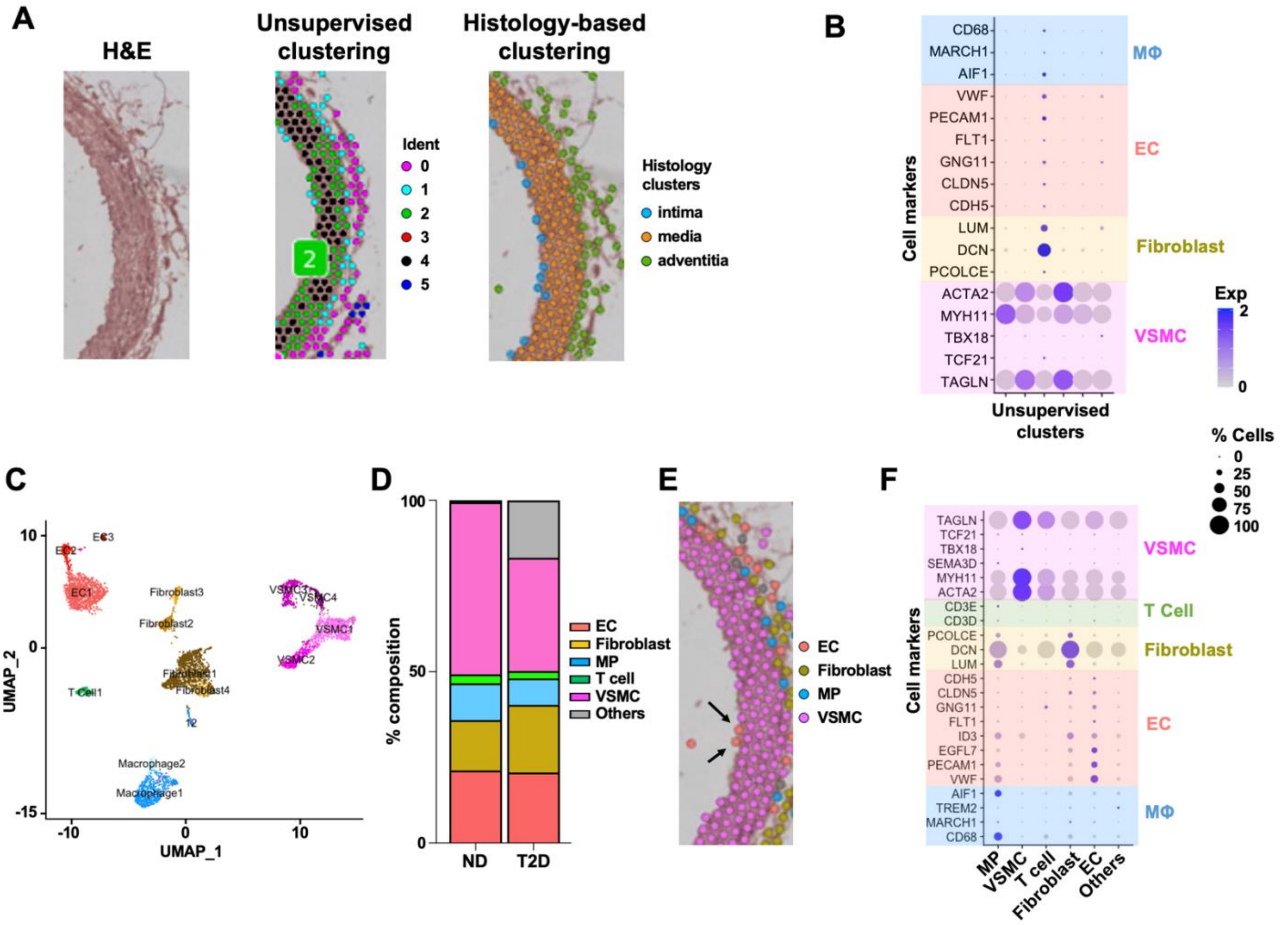


**Fig. S13. Integration of ST- and scRNA-seq. (A)** Representative H&E staining of the mesenteric arterial wall (left) and the superimposed unsupervised clustering (center) and histology-based clustering of ST-seq data (right). (**B)** Expression of cell marker genes (y-axis) of unsupervised clustering analysis of ST-seq. Note the expression of VSMC markers in all clusters and that of marker gene representative of multiple cell types in some cluster. **(C,D)** Analysis of whole mesenteric artery scRNA-seq, with UMAP showing 5 major cell types captured (in C) and bargraph showing % composition of total cell population. **(E,F)** Integrative analyses of ST- and scRNA-seq. Representative annotation of each voxel in the ST-seq with the most dominant cell type, with arrows indicating EC-dominant voxels in the intima (in E). Dotplot showing expression of marker genes in annotated clusters (in F). Note the expression of specific cell type markers (labeled on y-axis) in voxels with the corresponding dominant cell types (labeled on x-axis).

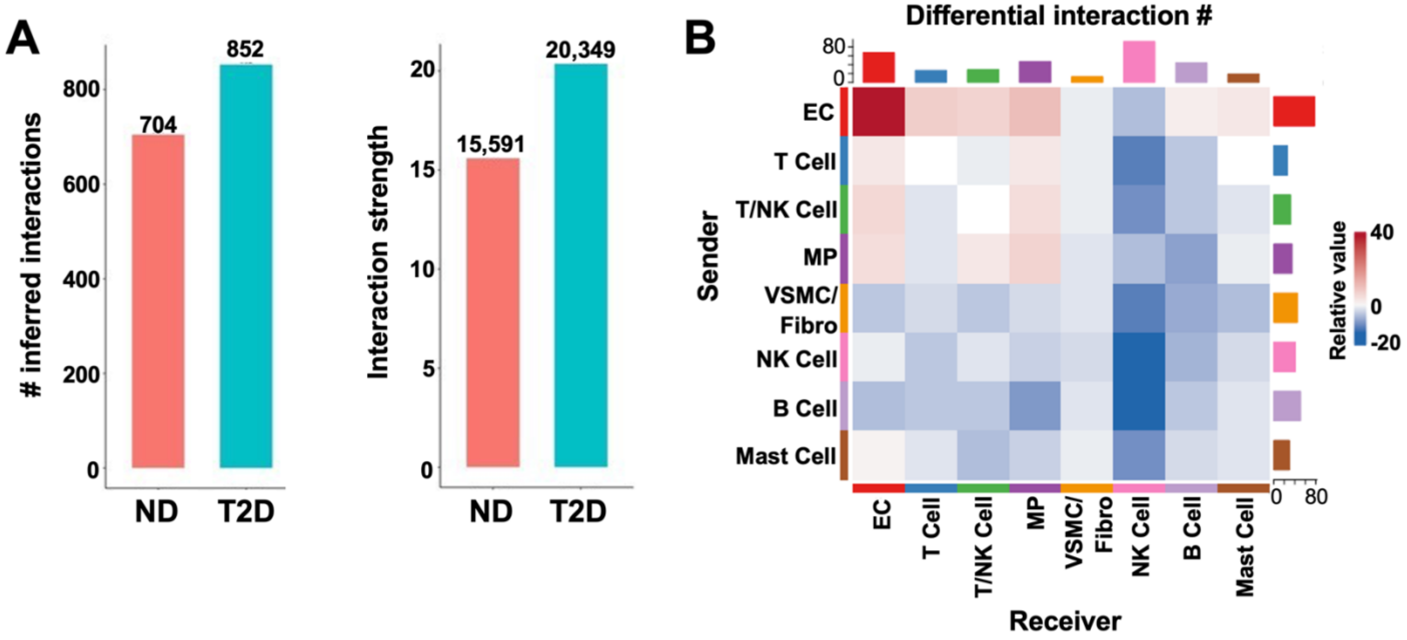


**Fig. S14. Cell communication network in the mesenteric artery intima. (A)** Bar chart plotted with number (#) and strength of the inferred cell-cell interactions identified from intima scRNA-seq. **(B)** Heatmap showing differential numbers of interactions between individual cell types associated with T2D as compared to ND. Gradient denotes relative values, with red indicating up-regulated and blue indicating down-regulated.

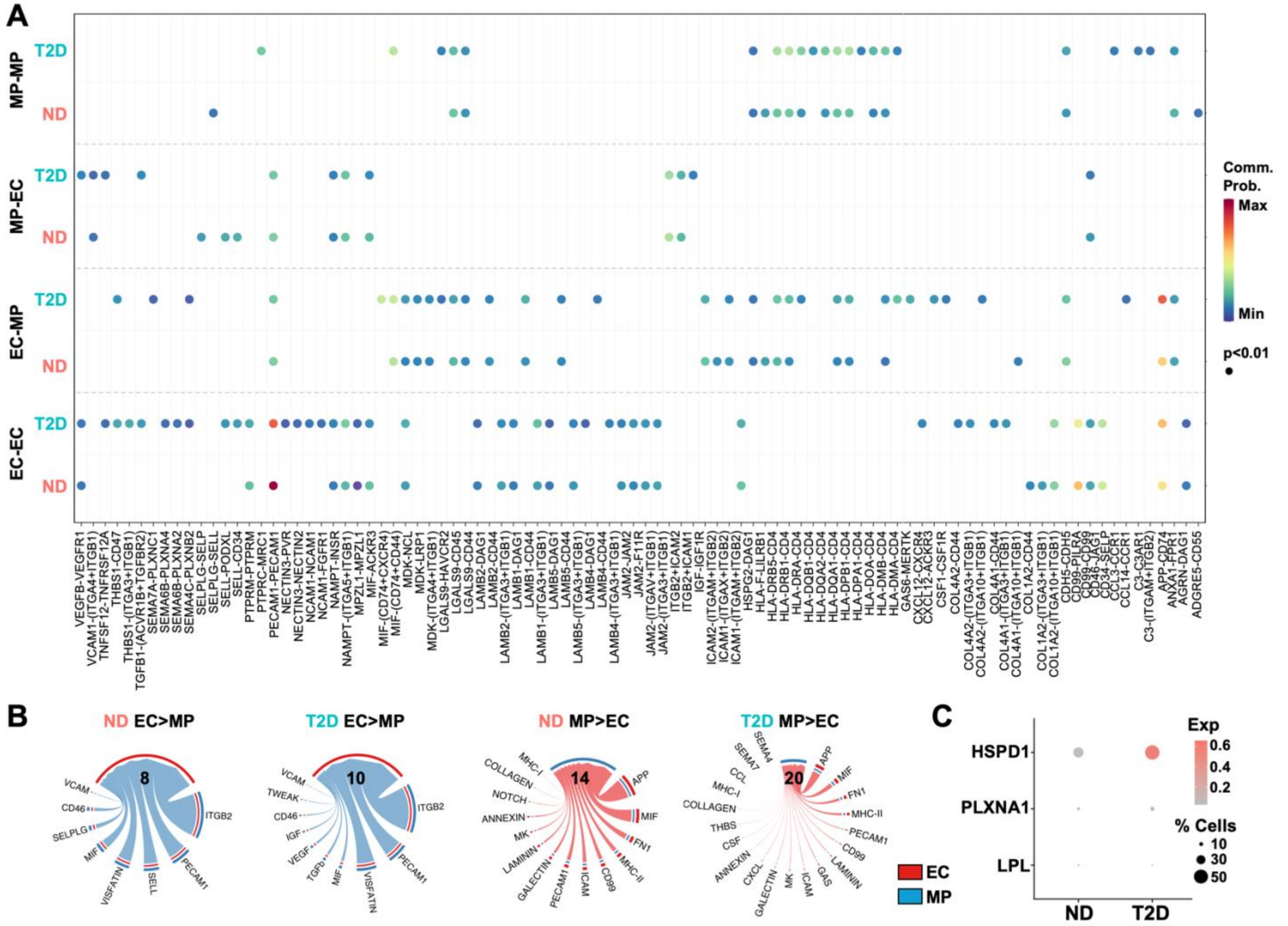

**Fig. S15. EC-MP communication network. (A)** Dotplot of significant receptor-ligand complexes formed between EC and EC, EC and MP, MP and EC and MP and MP across ND and T2D. Scale bar indicates communication probability. **(B)** Chord diagram of significant pathways initiated from ECs to MP (left) and from MP to ECs (right) in ND and T2D. The outer ring shows the sender and receiver cells, with red indicating ECs and blue indicating MPs. The values in the diagrams denote the number of pathways between the communicating cells. Note the increase in the number of bi-directional EC-MP interactions in T2D. **(C)** Expression of additional TREM2 ligands in ECs revealed by intima scRNA-seq.

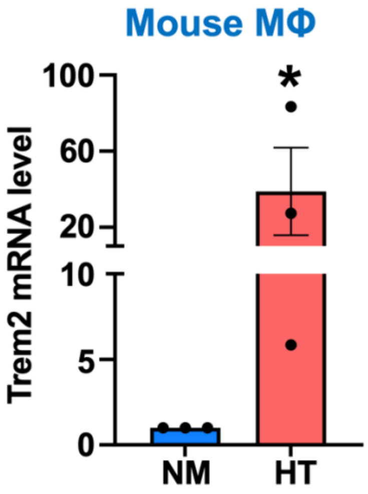


**Fig. S16. Induction of TREM2 in HT-treated mouse MΦ.** qPCR of *Trem*2 in mouse MΦ (Raw 264.7) under NM or HT. Graphs represent mean±SEM from 3 independent experiments and *$p<0.05$ based on Student's t-test.

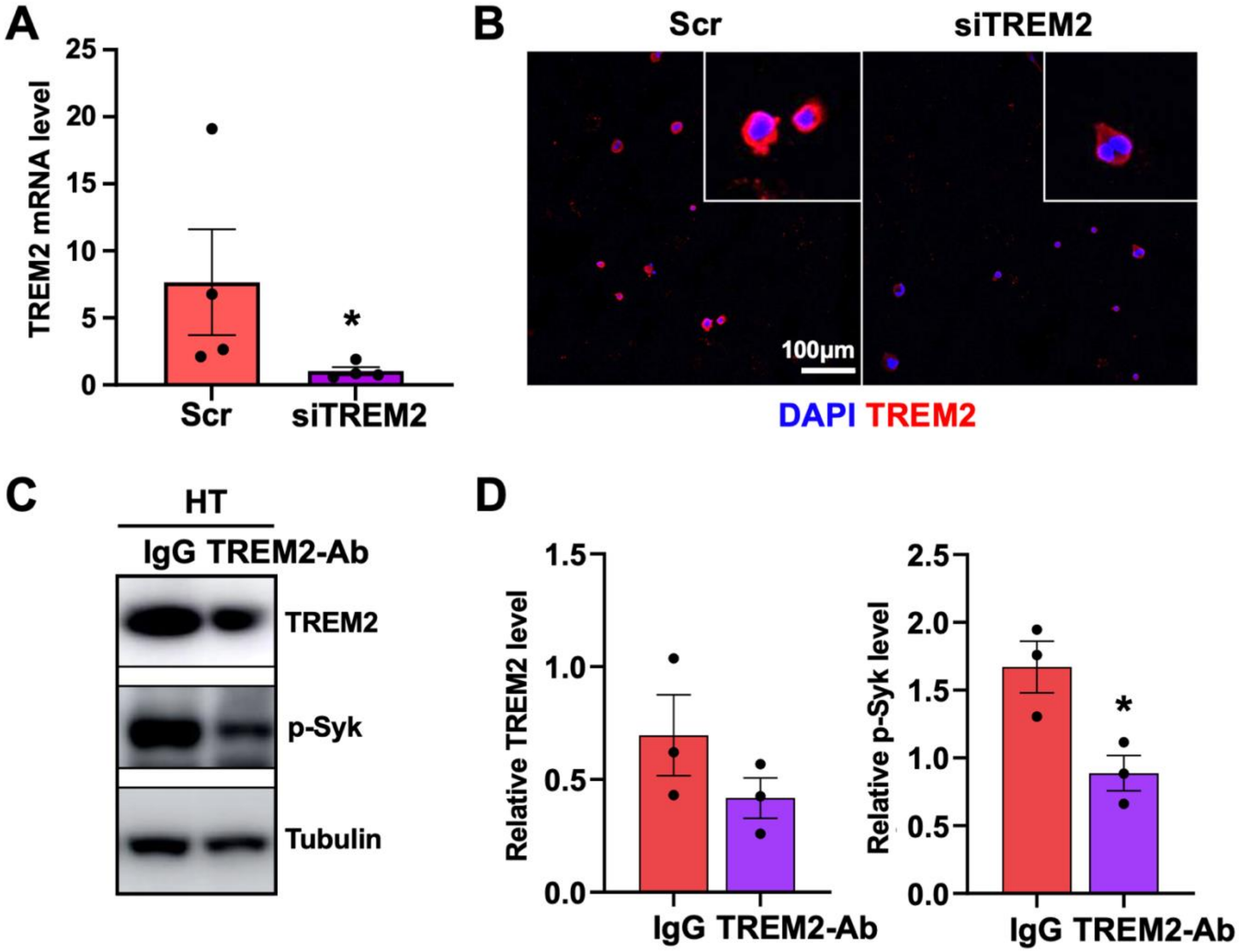


**Fig. S17. Effect of TREM2 siRNA and TREM2-Ab. (A)** TREM2 mRNA levels in THP-1-derived MΦ transfected with scramble (Scr) of TREM2 siRNA (siTREM2) and treated with HT. **(B)** Immunofluorescence of TREM2 with DAPI counterstain in the same experiment as in (A), with Zoom-in images. **(C, D)** Immunoblotting of TREM2 and p-Syk in THP-1-derived MΦ treated with TREM2-Ab or IgG control. Representative immunoblots (in C) and quantification of TREM2 and p-Syk, normalized to tubulin (in D). Data represent mean±SEM from 3-4 independent experiments. *$p<0.05$ based on Student's t-test.

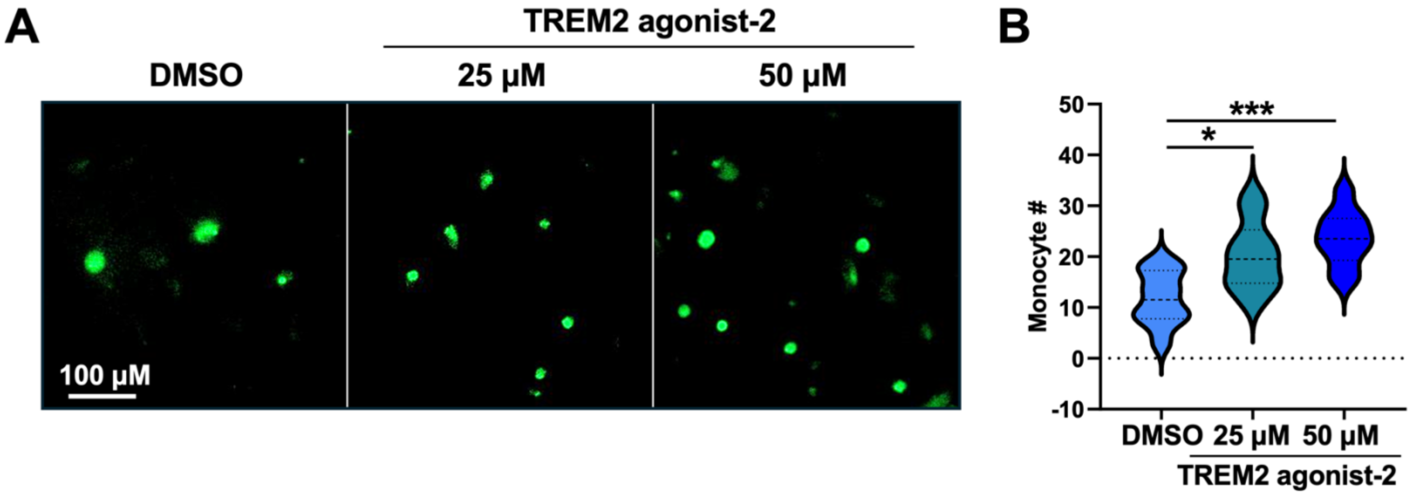


**Fig. S18 Effect of TREM2 agonist on monocyte-EC adhesion. (A) Representative images from** monocytes pre-treated with DMSO or TREM2 agonist-2 (25 or 50 μM) for 16 hours, labeled by CMFDA, and incubated with NM-treated HUVECs. **(B)** Quantification of monocyte adhesion from 3 independent experiments. *$p<0.05$, ***$p<0.001$ based on one-way ANOVA followed by Dunnett's test.

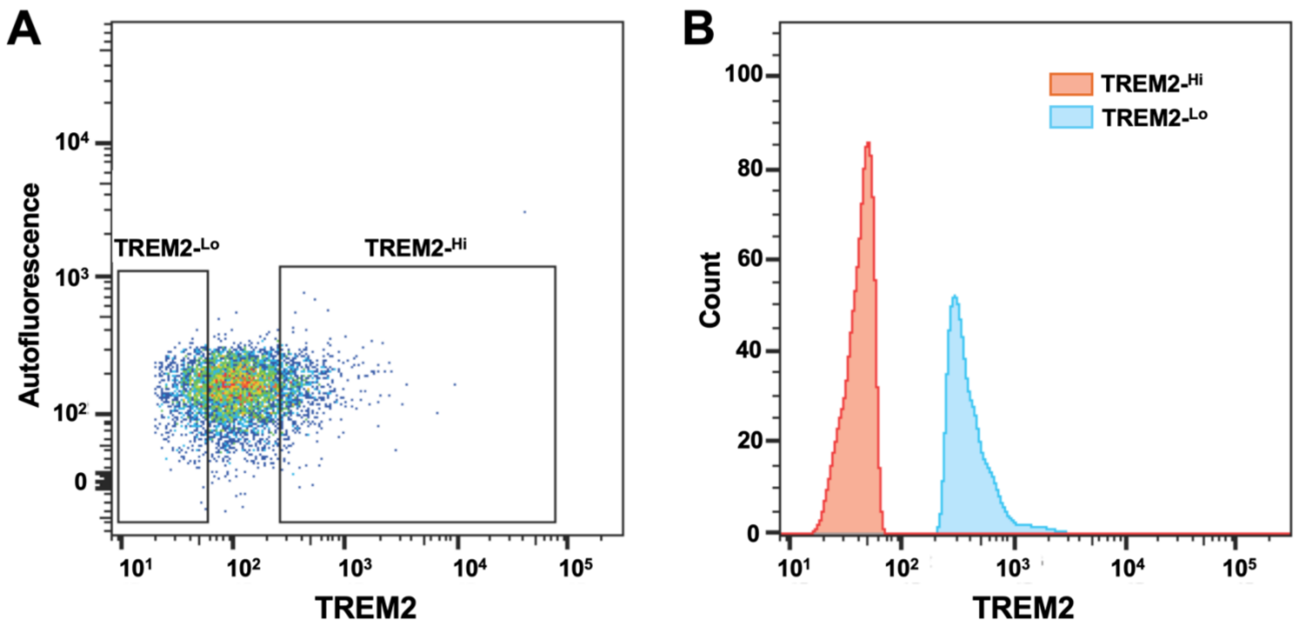


**Fig. S19. FACS of TREM2-high (Hi) and TREM2-low (Lo) cells from human spleen. (A)** Gating strategy for FACS of TREM2$^{-Hi}$ and TREM2$^{-Lo}$ cells from human spleen from donors with T2D. **(B)** Histogram showing separation of TREM2$^{-Hi}$ and TREM2$^{-Lo}$ cells based on TREM2 expression**.**

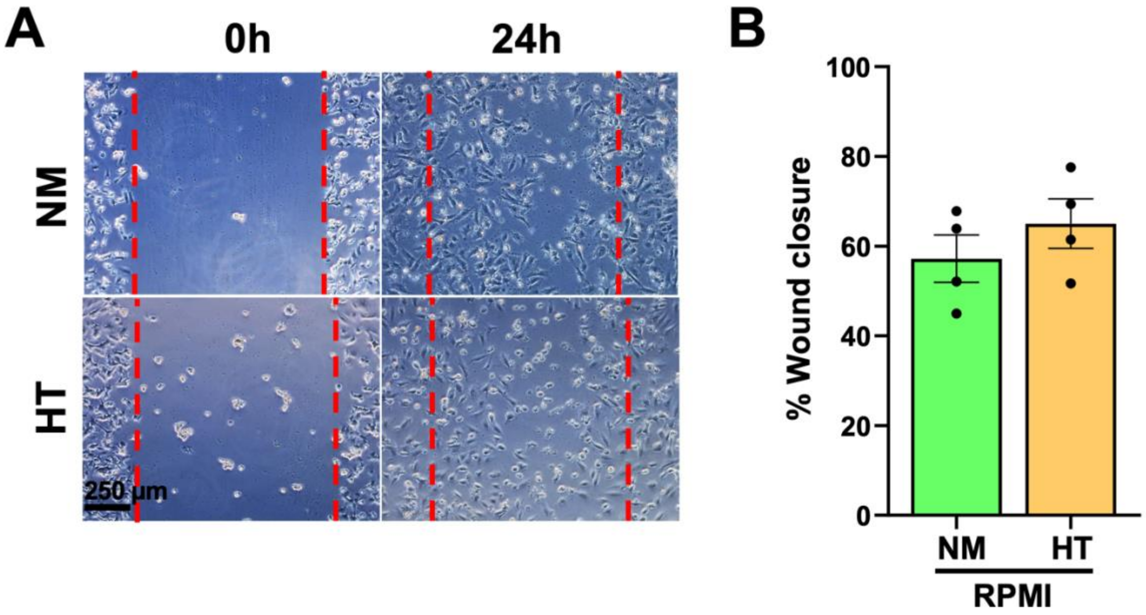


**Fig. S20. Effect of culture media on EC migration.** Representative images (in A) and quantification (in B) of scratch wound assay using ECs incubated with RPMI supplemented with 25mM mannitol (NM) or 25 mM glucose plus TNF-$\alpha$ (HT) for 24 hours. Data represent mean±SEM from 4 independent experiments.

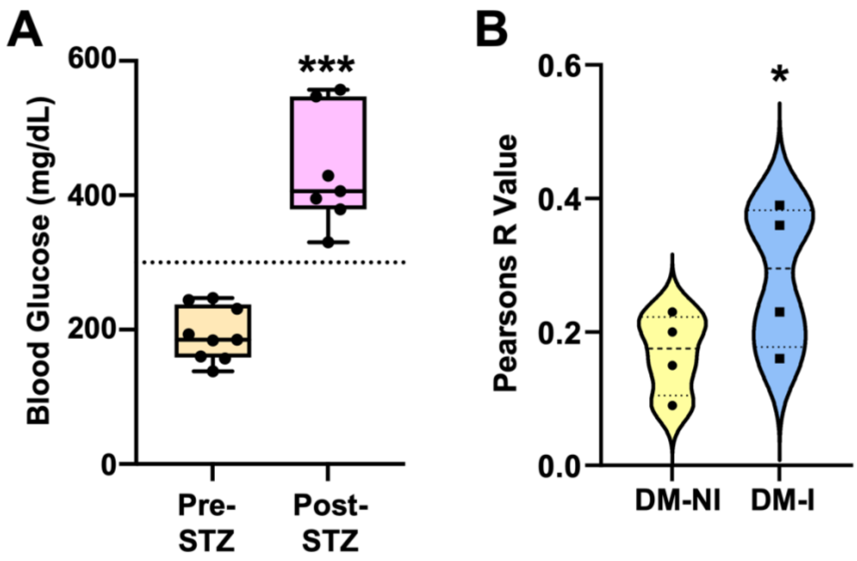


**Fig. S21. Characterization of the mouse model with DM-HLI. (A)** Confirmation of hyperglycemia (glucose >300 mg/dL) post STZ injections. **(B)** Quantification of TREM2 and IB4 co-IF staining on NI vs I limbs. *p<0.05, ***p<0.001 based on Student's t-test.

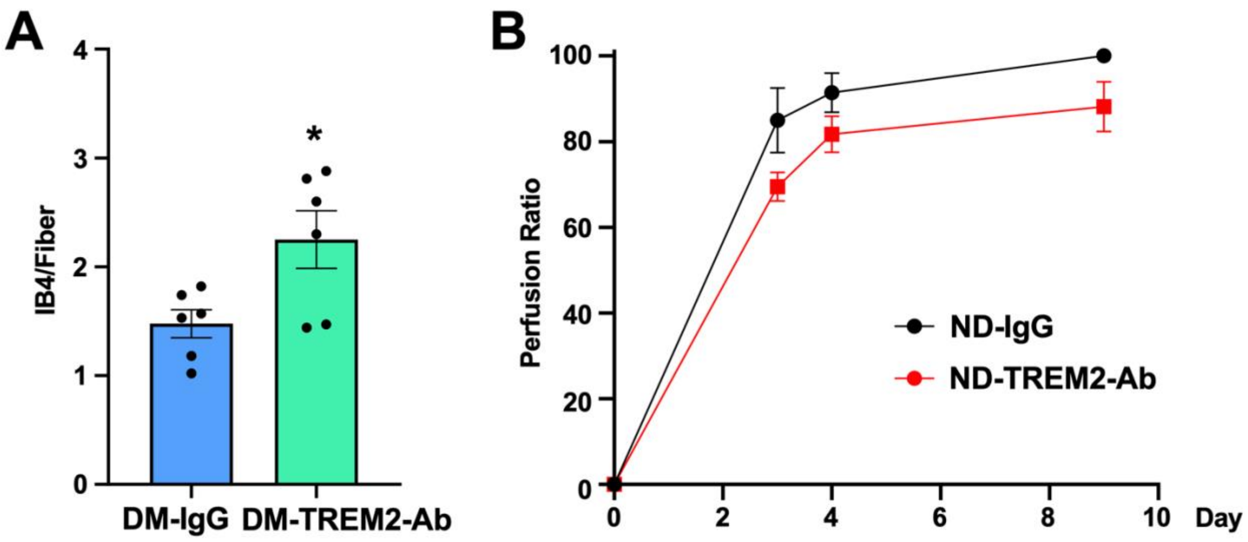


**Fig. S22. Effect of TREM2-Ab on mice with HLI. (A)** Quantification of IB4 staining of ischemic limb of STZ-treated (i.e., DM) mice receiving IgG or TREM2-Ab (n=6/group). **(B)** Quantification of flow perfusion in ND mice receiving IgG or TREM2-Ab (n=5/group). **p<0.01 based on Student's t-test.

### Table S1. Donor characteristics for human mesenteric artery profiling and cell isolation

| Donor | Age | Sex | Ethnicity | BMI | HbA1c (%) | Medical History |
|---|---|---|---|---|---|---|
| ***Intima scRNA-seq*** | | | | | | |
| ND-1 | 20 | M | Hispanic | 26 | 5.5 | No T2D-associated co-morbidities |
| ND-2 | 47 | M | Caucasian | 26.3 | 5.5 | No T2D-associated co-morbidities |
| ND-3 | 62 | M | Caucasian | 26 | 5.1 | Hypertension and high cholesterol for 5 yrs |
| ND-4 | 50 | M | Caucasian | 38.6 | 5.8 | Hypertension and hyperlipidemia for 1 yr |
| ND-5 | 59 | M | Caucasian | 28 | 5.4 | Hypertension for unknown duration |
| T2D-1 | 42 | F | Hispanic | 40 | 9.5 | Untreated T2D 25 yrs, hypertension for >10 yrs, family history of DM |
| T2D-2 | 46 | M | Hispanic | 45.8 | 6.9 | Untreated T2D for unknown duration, high cholesterol, family history of DM |
| T2D-3 | 63 | M | Hispanic | 35.9 | 7 | Untreated T2D for 20 yrs, hypertension and hyperlipidemia for 20 yrs, family history of DM |
| T2D-4 | 56 | M | Hispanic | 27.7 | 8.5 | Untreated T2D for 7 yrs, hypertension for 40 yrs. |
| T2D-5 | 63 | M | Caucasian | 33.9 | 6.7 | Treated T2D for 10 yrs but non-compliant, hypertension for 10 yrs, and family history of DM |
| ***ST-seq*** | | | | | | |
| ND-6 | 62 | M | Caucasian | 26 | 5.1 | Hypertension and hypercholesterolemia for 5 yrs |
| ND-7 | 44 | M | Caucasian | 24.4 | 5.3 | No T2D-associated co-morbidities |
| T2D-6 | 34 | M | Hispanic | 27.6 | 7.5 | Treated T2D for 7 yrs, hypercholesterolemia for 5 yrs |
| T2D-7 | 58 | M | Hispanic | 22.9 | 10.5 | Treated T2D for 8 yrs, treated hypertension for 15 yrs |
| ***Whole artery scRNA-seq*** | | | | | | |
| ND-8 | 62 | F | African American | 26.8 | 6.2 | Treated hypertension for > 5 yrs |
| ND-9 | 25 | M | Hispanic | 23.7 | 4.9 | Family history of DM |
| T2D-8 | 57 | F | Asian | 30.00 | 6 | Treated T2D for 1 yr, hypertension and hypercholesterolemia for 3 yrs |
| ***IHC staining*** | | | | | | |
| ND-10 | 29 | F | Hispanic | 24.1 | 4.3 | No T2D-associated co-morbidities |
| ND-11 | 18 | M | Hispanic | 20.1 | 4.9 | No T2D-associated co-morbidities |
| ND-12 | 57 | F | Caucasian | 24 | 4.4 | Hypercholesterolemia for unknown duration |
| T2D-9 | 56 | M | Hispanic | 29 | 7.6 | Treated DM > 10 yrs, hypertension for unknown duration |
| T2D-10 | 50 | F | Caucasian | 35 | 9.9 | Treated T2D for 30 yrs but non-compliant. Family history of DM |
| T2D-11 | 58 | F | Caucasian | 35 | 10.1 | Unknown |
| ***FACS-based cell isolation from spleen*** | | | | | | |
| T2D-12 | 63 | M | Caucasian | 32.1 | 6.5 | No T2D-associated co-morbidities |
| T2D-13 | 53 | M | Caucasian | 28.7 | 6.3 | Treated T2D for 1 yr, treated hypertension for 7 yrs |

### Table S2: Cell and cluster composition (%) of intima scRNA-seq

| Sample | EC1 | EC2 | MP-C1 | MP-C2 | MP-C3 | MP-C4 | MP-C5 | T/NK1 | T/NK2 | T/NK3 | T/NK4 | T/NK5 | VSMC1 | VSMC2 | B Cell1 | B Cell2 | Mast Cell |
|---|---|---|---|---|---|---|---|---|---|---|---|---|---|---|---|---|---|
| ND | 24.0 | 8.9 | 3.4 | 3.2 | 3.0 | 4.3 | 2.4 | 10.5 | 2.9 | 6.4 | 1.0 | 5.9 | 19.3 | 1.0 | 3.6 | 0.1 | 0.2 |
| T2D | 21.4 | 5.2 | 12.2 | 7.8 | 6.1 | 3.6 | 0.3 | 12.1 | 2.5 | 12.3 | 2.3 | 5.5 | 4.6 | 1.0 | 1.3 | 0.9 | 0.9 |

**Table S3. Top 50 up- and down-regulated genes in ECs identified by intima scRNA-seq.**

| Gene | avg_log2FC |
|---|---|
| NOS3 | -0.88222 |
| MT2A | -1.342594 |
| FGFR3 | -0.094289 |
| IFI6 | -1.379431 |
| AP2B1 | -0.341866 |
| ZBTB16 | -0.256731 |
| SLC12A6 | -0.265977 |
| DMAC2 | -0.235564 |
| CKB | -0.758243 |
| CBFA2T3 | -0.232218 |
| PLAT | -1.054735 |
| GIMAP6 | -0.183532 |
| ST6GALNAC1 | -0.414558 |
| GPX3 | -1.56035 |
| AC097534.2 | -0.149144 |
| TRIB2 | -0.418396 |
| DOCK9 | -0.575661 |
| NKTR | -0.491195 |
| ODF2L | -0.192217 |
| LAMA5 | -0.427778 |
| COPS9 | -0.458208 |
| NES | -0.730237 |
| NRGN | -0.382156 |
| ABCB1 | -0.078795 |
| ADAM15 | -1.085672 |
| KDM5D | -0.276549 |
| NAPRT | -0.27277 |
| RPS27L | -0.346113 |
| SNCAIP | -0.385851 |
| SEC62 | -0.193422 |
| SORBS2 | -0.306715 |
| CLIC2 | -0.209801 |
| IFITM2 | -0.19717 |
| LRP5 | -0.245313 |
| EMP2 | -0.450262 |
| NPR1 | -0.667113 |
| TTTY14 | -0.403315 |
| ST8SIA6 | -0.533657 |
| TRIM25 | -0.297438 |
| MXD4 | -0.455658 |
| MMRN2 | -0.489797 |

| Gene | avg_log2FC |
|---|---|
| PECAM1 | -0.668769 |
| LENG8 | -0.298186 |
| NRP1 | -0.270124 |
| PTPRN2 | -0.138284 |
| PLA1A | -0.644795 |
| HSPA12B | -0.595726 |
| PTGIS | -0.792855 |
| CADPS2 | -0.353209 |
| CCND1 | -0.679067 |
| BTG1 | 0.321091 |
| TMED7 | 0.276233 |
| TXNRD2 | 0.804899 |
| SLPI | 1.791946 |
| DUSP6 | 0.558254 |
| DEPP1 | 1.153911 |
| HAPLN3 | 0.725136 |
| FN1 | 1.66825 |
| CCN2 | 2.09406 |
| FARP1 | 0.401291 |
| AL021155.5 | 0.238298 |
| MAP3K8 | 0.294974 |
| THEM6 | 0.496491 |
| MRPS6 | 0.33715 |
| EIF3L | 0.329593 |
| TMEM98 | 0.546909 |
| EEF1B2 | 0.595924 |
| RPS3 | 0.35229 |
| HIST1H1E | 0.28656 |
| PLEKHO1 | 0.275106 |
| ATP6V1G1 | 0.381715 |
| EEF1A1 | 0.38929 |
| TAF9 | 0.125919 |
| THBS1 | 0.99706 |
| SPNS2 | 0.333826 |
| IER3 | 2.267887 |
| FNBP1 | 0.263441 |
| PLAAT3 | 0.278749 |
| PPP1R15A | 1.201146 |
| FAM43A | 1.640964 |
| MRPL51 | 0.173223 |
| SEMA6B | 0.205945 |

| Gene | avg_log2FC |
|---|---|
| BEX4 | 0.444303 |
| FBLIM1 | 0.673932 |
| ANP32A | 0.428096 |
| NFKBIZ | 0.297763 |
| BNIP3L | 0.394932 |
| MTUS1 | 0.582252 |
| SRSF7 | 1.36425 |
| MFAP2 | 0.620265 |
| IGFBP7 | 1.021734 |
| FEZ2 | 0.367289 |
| STEAP2 | 0.484391 |
| MBOAT7 | 0.169991 |
| PAWR | 0.513839 |
| AC090152.1 | 0.135176 |
| AC104083.1 | 0.378193 |
| HTR4 | 0.254794 |
| CARD19 | 0.195888 |
| RPL3 | 0.611142 |

**Table S4. CLTI patient characteristics**

| Donor | Age | Sex | Ethnicity | BMI | DM* | Side | Surgery |
|---|---|---|---|---|---|---|---|
| PAD-1 | 87 | M | African American | 24.8 | 0 | R | Above-Knee Amputation |
| PAD-2 | 59 | M | American Indian | 35.3 | 1 | R | Above-Knee Amputation |
| PAD-3 | 84 | M | Caucasian | 30.4 | 1 | L | Below-Knee Amputation |
| PAD-4 | 72 | M | Caucasian | 38.4 | 1 | L | Below-Knee Amputation |
| PAD-5 | 83 | M | Caucasian | 27.5 | 1 | R | Above-Knee Amputation |
| PAD-6 | 83 | M | Caucasian | 27.5 | 1 | L | Above-Knee Amputation |
| PAD-7 | 71 | F | Caucasian | 36.4 | 1 | L | Above-Knee Amputation |
| PAD-8 | 72 | M | Caucasian | 23.1 | 1 | R | Below-Knee Amputation |

* 0=ND and 1=DM

**Table S5. Primers and siRNA sequences**

| Gene/ID | Species | Sequence |
|---|---|---|
| **TREM2** | Human | Forward CTGCTCATCTTACTCTTTGTCAC<br>Reverse CAGTGCTTCATGGAGTCATAGG |
| **ICAM1** | Human | Forward GTGTCCTGTATGGCCCCCGACT<br>Reverse ACCTTGCGGGTGACCTCCCC |
| **VCAM1** | Human | Forward TTTTCGGAGCAGGAAAGCCC<br>Reverse GTCAATGTTGCCCCCAGAGA |
| **ACTB** | Human | Forward CATGTACGTTGCTATCCAGGC<br>Reverse CTCCTTAATGTCACGCACGAT |
| **Trem2** | Mouse | Forward TTGCTGGAACCGTCACCATC<br>Reverse CACTTGGGCACCCTCGAAAC |
| **36B4** | Mouse | Forward AGATTCGGGATATGCTGTTGGC<br>Reverse TCGGGTCCTAGACCAGTGTTC |
| **ON-TARGETplus SMARTpool TREM2 siRNA** | Human | 1. AGAGACACGUGAAGGAAGA<br>2. GGACACAUCCACCCAGUGA<br>3. GGUAUCAGCUCCAAACUCU<br>4. GGUCAGCACGCACAACUUG |
